# How a pathogenic mutation impairs Hsp60 functional dynamics from monomeric to fully assembled states

**DOI:** 10.1101/2024.09.09.611948

**Authors:** Luca Torielli, Federica Guarra, Hao Shao, Jason E. Gestwicki, Stefano A. Serapian, Giorgio Colombo

## Abstract

Heat Shock Protein 60 kDa (Hsp60) is a mitochondrial chaperonin that cooperates with Hsp10 to drive the correct folding of client proteins. Monomers **M** of Hsp60 (featuring equatorial, intermediate, and apical domains) first assemble into 7-meric Single rings (**S**), then pairs of **S** interface equatorially to form 14-meric Double rings (**D**) that accommodate clients into their lumen. Recruitment of 7 Hsp10 molecules per pole turns **D** into a 28-meric Football-shaped complex (**F**). Sequential hydrolysis of ATP present in each Hsp60 unit of **F** finally drives client folding and **F** disassembly. Equatorial domain mutation V72I occurs in SPG13, a form of hereditary spastic paraplegia: while distal to the active site, this severely impairs the chaperone cycle and stability. To understand the molecular bases of this impairment we have run atomistic molecular dynamics (MD) simulations of **M**, **S**, **D**, and **F** for both WT and mutant Hsp60, with two catalytically relevant Hsp60 aspartates in **D** and **F** modelled in three different protonation states. Additionally, **D** in one protonation state was modelled post-hydrolysis (total production time: 36 µs). By combining complementary experimental and computational approaches for the analysis of functional dynamics and allosteric mechanisms, we consistently find that mutation V72I significantly rewires allosteric routes present in WT Hsp60 across its complexes, from isolated **M** units right up to **F**, rigidifying them—as observed experimentally—by introducing a direct allosteric link between equatorial and apical Hsp60 domains that bypasses the ATP binding site (wherein we observe the alteration of mechanisms driving reactivity). Our results reveal a multiscale complexity of functional mechanisms for Hsp60 and its pathogenic mutant, and may lay the foundation for the design of experiments to fully understand both variants.

## Introduction

The human mitochondrial Heat shock protein weighing 60 kDa (Hsp60) is a molecular chaperone that belongs to Group I chaperonins, a family of biomolecules fundamental for the control of proteostasis under both normal and stress conditions.^[1]^ Alterations in Hsp60 homeostasis and function have been related to pathological conditions such as cancers,^[2]^ diabetes and Alzheimer’s disease,^[3]^ and this has driven interest in understanding more about the structure and function of this chaperone system. Generally, group I chaperonins are ATPases^[4]^ that assemble into single or double heptameric ring complexes, assisting in the folding of denatured or nascent proteins within a discrete folding chamber.^[5]^ Each monomer (**M**) of Hsp60 contains an equatorial, apical, and intermediate domain (Figure 1a, left).^[6]^ The equatorial domain includes the ATP binding site (Figure 1b) and it is also involved in multiple inter- and intra-ring interactions to stabilise oligomers. A C-terminal region of the equatorial domain is also involved in binding to client proteins. The apical domain is responsible for interactions with the co-chaperone Hsp10 and client proteins. Finally, the intermediate domain includes the hinge region that controls inter-domain movements, and it bears helix α14 (Figure 1c; *vide infra*) that contains catalytic residue Asp397.^[7]^ This architecture allows Hsp60 to couple ATP binding and hydrolysis to many important conformational changes, both within the protomers and across its oligomeric complexes. Yet, there are many important questions remaining about the molecular mechanism of Hsp60 functions.

**Figure 1.**
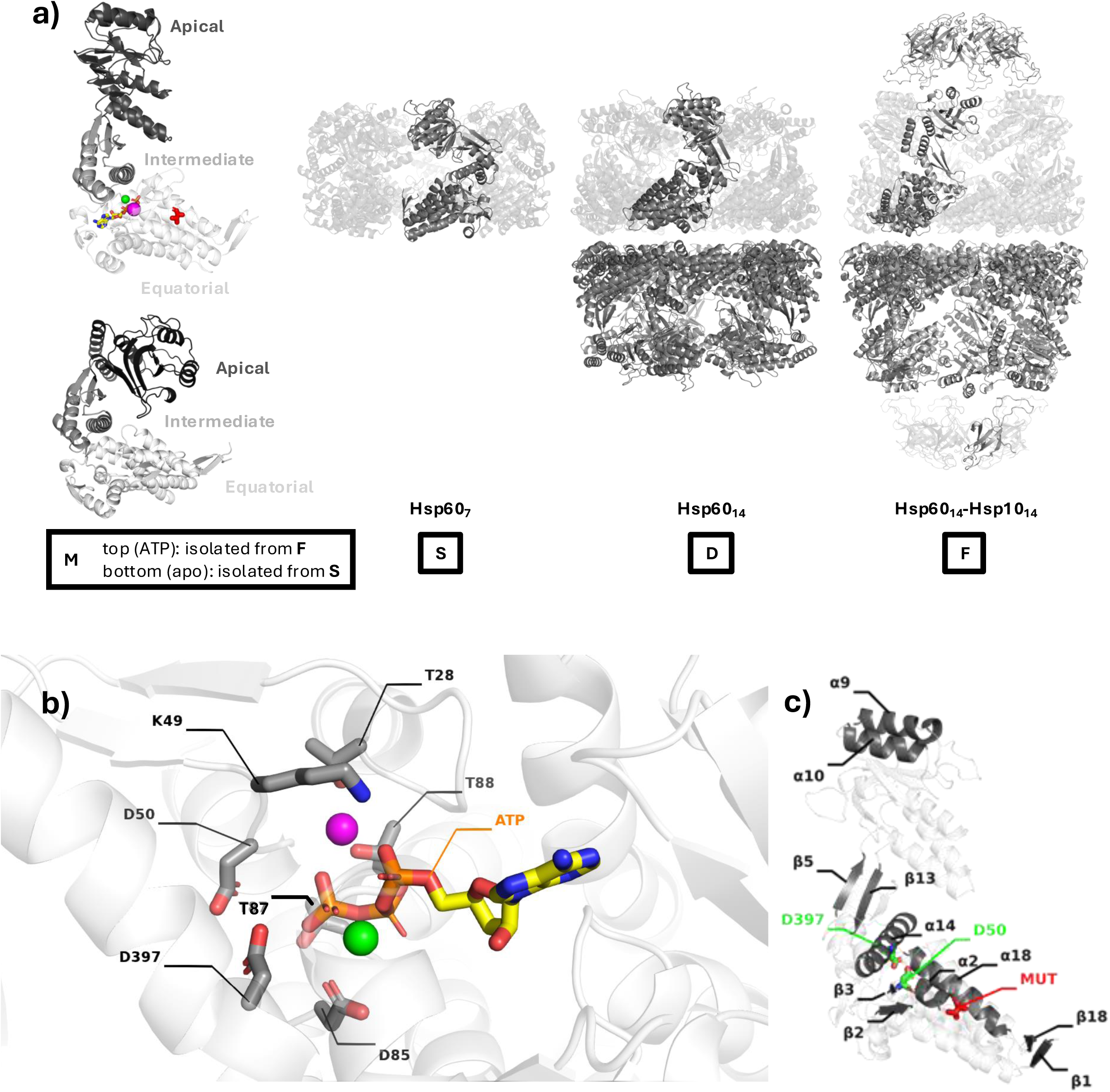
Structural aspects of Hsp60 complexes. a) Hsp60 monomer **M** and higher Hsp60[-Hsp10] complexes simulated in this work, as prepared from their CryoEM structures (see main text for details). Two instances of **M** are represented on the left: the bottom (apo) instance, isolated from single ring **S**, is the one featured in our simulations. In the top instance, isolated from football-shaped complex **F** and not featured in our simulations, we have marked salient elements, including the mutating Val72 as red sticks; ATP (C: yellow sticks; N: blue sticks; P: orange sticks; O: red sticks); Mg^2+^ (green sphere); K^+^ (magenta sphere). Both instances of **M** have their equatorial, intermediate, and apical domains clearly marked. In all systems, hydrogen, Na^+^, Cl^−^, waters are omitted for clarity. A number of Hsp60 and Hsp10 monomers in higher complexes are shown nontransparently, for reference. b) Representation of the active site with ATP and salient residue side-chains marked as sticks with conventional element colours (except for ATP carbons, in yellow). Mg^2+^ and K^+^ cations are shown as spheres (green and magenta, respectively). c) Salient secondary structure components of the Hsp60 monomers, as mentioned in the text, illustrated on a monomer isolated from the football-shaped complex. For reference, we retain stick representations of ATP, of catalytic aspartates Asp50 and Asp397 (C atoms in green), and of the mutation site (all atoms in red). Cations Mg^2+^ and K^+^ are represented as green and magenta spheres, respectively.

The homologous bacterial system, GroEL/GroES, has been the subject of long-standing research to understand chaperonin structure-function.^[8]^ That work has often been used to make assumptions about how human Hsp60/Hsp10 works. Indeed, more recent structural and biochemical investigations have shown that the functional cycle of Hsp60/Hsp10 chaperonin/cochaperonin shares conserved intermediate states with that of GroEL/GroES.^[9]^ However, important differences have also been observed. For example, the stability of Hsp60/10 oligomeric assemblies and inter-ring cooperativity are quite different from GroEL/GroES.^[10]^ Indeed, the relatively poor stability of Hsp60 complexes *in vitro* has limited mechanistic investigations until recently.^[11]^ While GroES readily assembles into tetradecameric double rings even in absence of ATP,^[12]^ intra- and inter-ring interactions existing in Hsp60 are weaker,^[9d]^ with the apo protein shown to exist in a dynamic equilibrium between single and double-ring complexes or even to dissociate into monomers.^[11, 13]^ Both apo and ATP-bound Hsp60 complexes were found to be competent client binders.^[14]^ With no evidence for inter-ring negative cooperativity,^[9]^ it is ATP binding that shifts the equilibrium back in favour of double-ring complexes **D** with two heptameric single rings **S** stacked back-to-back (Figure 1a).^[11, 13]^ This is also because ATP binding induces significant conformational changes which entail the closure of intermediate helix α14 and its Asp397 (Figure 1b, Figure 1c) and elevation of the apical domain from its “closed”, lowered position in the apo form. In the GroEL/GroES system, the latter event is associated with the progressive decrease of affinity for the client and increase in affinity for cochaperonins.^[15]^ Finally, assembly of the two heptameric Hsp10 caps (7 Hsp10 units at each apex) leads to the formation of the American-football-shaped structure **F** (Figure 1a): this is the complex that contains a fully functional folding chamber, wherein client protein folding can occur in a sealed, protected environment. Following ATP hydrolysis, the folding chamber disassembles enabling the release of folded clients or partially folded intermediates.^[16]^ Unlike GroEL/GroES, half-football structures formed by a heptameric Hsp60 ring and a heptameric Hsp10 cap were also observed and are regarded as having functional relevance.^[9a, 9b, 16]^ The accepted view is that ATP hydrolysis is featured by intra-ring positive cooperativity with monomers in the ring(s) working concertedly.^[17]^ Interestingly, recently solved client-bound structures of Hsp60/Hsp10 have given evidence for asymmetric ‘up-down’ arrangements of the apical domains in apo single-rings and ATP-bound double rings.^[14]^ These conformations allow efficient client retention while cochaperonin units are recruited and mechanistically explain the observed overlap in client/cochaperonin-interacting regions on Hsp60. Symmetry is re-established in the Hsp60/Hsp10 complex once the stabilisation of the ‘up’ conformations by Hsp10 enlarges the chamber volume for client folding. Despite these major advances in recent years, there are still important questions remaining to be addressed. Namely, while the “snapshots” of Hsp60/Hsp10 structure have suggested potential molecular mechanisms of allostery, the intermediates along these trajectories have remained obscured.

We saw a potential way to address these questions by combining molecular dynamics (MD) with a disease- associated point mutation in Hsp60. Point mutations in Hsp60’s equatorial domain have been identified as responsible for neurodegenerative diseases.^[18]^ Among these, mutation V72I (numbering refers to the mature protein) causes hereditary spastic paraplegia SPG13, characterised by progressive weakness and spasticity of the lower limbs.^[18d]^ Because this mutation has such a strong phenotype in humans, it suggests that it has an important impact on function. Yet, V72I is not in contact with the nucleotide or the active site, so its effects are likely to be allosteric. This is quite surprising, because the mutant Isoleucine side chain is only one methylene group longer than the wild type Valine. *In vitro* experiments have shown that V72I stabilises the Hsp60 oligomer, while reducing client folding activity and impacting ATP hydrolysis.^[14, 19]^ Thus, we reasoned that an understanding of the mechanistic determinants of these effects would lead to a deeper understanding of Hsp60 function. For example, it is not clear whether the increase in ATPase activity is due to an intrinsic increase in the enzymatic reaction rate or an apparent effect due to a combination of increased oligomerisation and cooperativity.^[14, 19–20]^ A recent computational study on the apo single ring focused on the mechanisms of increased oligomer stability in the V72I mutant; however, mechanistic details on how this change reduces folding capacity are still lacking.^[19–20]^ In addition, it is not clear how V72I mutations have an impact on mixed oligomers, containing both mutant and WT protomers. This is an important question because the mutation is dominant and effective in the heterozygous state involving mixed V72I and WT protomers.

Here, with a significant computational effort, we employ unbiased atomistic molecular dynamics simulations to unravel the dynamics of WT and V72I Hsp60 at multiple scales, encompassing most of the possible oligomeric organisation states of the protein and its assemblies (Figure 1a). We start from monomeric Hsp60 **M**, then move up one level to apo single ring **S**, then further to ATP-bound double ring **D**, and finally right up to football-shaped Hsp60_14_/Hsp10_14_ complex **F**. Through comparative analysis of the WT and V72I variants of **M** and its higher complexes—using different computational tools aimed at characterizing internal allosteric coordination patterns and pathways as well as overall conformational ensembles—we show that functional alterations in V72I Hsp60 can be reconnected to profound differences in its allosteric mechanisms compared to the WT variant. As a hallmark of the cooperative role of individual Hsp60 units in higher Hsp60 complexes, differences are detected in all complexes despite their origin from a deceivingly simple point mutation, and extend well beyond the modification of the reactivity of one single monomer.

Our simulations offer unique atomic-level insights into the effects of V72I mutation in terms of perturbation of functionally oriented internal dynamics, oligomeric stability, and asymmetric organisation of functional dynamic states; these insights are readily linkable to the impairment of folding activity, and could not have been obtained by simple structural comparison. For example, we provide the first evidence of the dynamic and structural consequences of combining wild-type *vs.* mutant monomers in oligomeric structures, reconnecting these consequences to important effects on the preorganisation of the system for client recognition and enzymatic reactivity. Overall, through our dynamic models of Hsp60 from single monomers to the largest complex of its functional cycle, not only do we precisely map the pathogenic impact of the V72I mutation, but we also provide important new findings on the dynamics and allostery of Hsp60 in general.

### Computational Methods

#### System Preparation

##### Starting structures

A total of 18 different mono- or multimeric systems are simulated in this work, as summarised in Table 1: 9 with WT Hsp60, and the same 9 with the corresponding V72I mutant. Throughout the article, we will adopt residue numbering for mature Hsp60: residues Met1-Tyr26 that constitute the mitochondrial import sequence are shed in mature Hsp60, and Ala27 becomes Ala1.

**Table 1.** Summary of the systems simulated in this work.

| Labels | Ligands | Hsp60 units | Hsp60 residues | Hsp10 units | Hsp10 residues | Asp50/Asp397 protonated |
| --- | --- | --- | --- | --- | --- | --- |
| **M**<sub>WT</sub> / **M**<sub>V72I</sub> | apo | 1 | 0–525 | 0 | – | No/No |
| **S**<sub>WT</sub> / **S**<sub>V72I</sub> | apo | 7 | 0–525 | 0 | – | No/No |
| **D**<sub>WT-AspAsp</sub> / **D**<sub>V72I-AspAsp</sub> | 14 × ATP/Mg<sup>2+</sup>/K<sup>+</sup> | 14 | 1–526 | 0 | – | No/No |
| **D**<sub>WT-AshAsp</sub> / **D**<sub>V72I-AshAsp</sub> | 14 × ATP/Mg<sup>2+</sup>/K<sup>+</sup> | 14 | 1–526 | 0 | – | Yes/No |
| **D**<sub>WT-AspAsh</sub> / **D**<sub>V72I-AspAsh</sub> | 14 × ATP/Mg<sup>2+</sup>/K<sup>+</sup> | 14 | 1–526 | 0 | – | No/Yes |
| **F**<sub>WT-AspAsp</sub> / **F**<sub>V72I-AspAsp</sub> | 14 × ATP/Mg<sup>2+</sup>/K<sup>+</sup> | 14 | 1–526 | 14 | 3–102 | No/No |
| **F**<sub>WT-AshAsp</sub> / **F**<sub>V72I-AshAsp</sub> | 14 × ATP/Mg<sup>2+</sup>/K<sup>+</sup> | 14 | 1–526 | 14 | 3–102 | Yes/No |
| **F**<sub>WT-AspAsh</sub> / **F**<sub>V72I-AspAsh</sub> | 14 × ATP/Mg<sup>2+</sup>/K<sup>+</sup> | 14 | 1–526 | 14 | 3–102 | No/Yes |
| **D**<sub>WT-ADP-Pi</sub> / **D**<sub>V72I-ADP-Pi</sub> | 14 × ADP/Pi/Mg<sup>2+</sup>/K<sup>+</sup> | 14 | 1–526 | 0 | – | No/Yes |

The nine systems simulated for each variant (WT, V72I) are (Table 1; Figure 1a): (1) the apo Hsp60 monomer **M**; (2) the apo single Hsp60_7_ ring **S**; (3–5) double Hsp60_14_ ring **D** in three different protonation states (*vide infra*) and with each Hsp60 unit ATP-, Mg^2+^-, and K^+^-bound; (6–8) Hsp60_14_/Hsp10_14_ football-shaped complex **F**, again in three different protonation states, and with each Hsp60 unit ATP-, Mg^2+^-, and K^+^-bound plus seven Hsp10 monomers capping each pole; and (9) again **D**, but this time with each Hsp60 unit ADP-, P_i_, Mg^2+^-, and K^+^-bound and only in one protonation state (with Asp397 protonated). These systems will be henceforth designated with general labels **M_WT_** / **M_V72I_**, **S_WT_** / **S_V72I_**, **D_WT_** / **D_V72I_**, and **F_WT_** / **F_V72I_**, **D_WT-ADP-Pi_** / **D_V72I-ADP-Pi_** respectively, regardless of protonation states.

Starting structures for **S_V72I_**, **D_V72I_**, **F_V72I_**, and **D_V72I-ADP-Pi_** were directly derived from CryoEM structures that have recently also been released into the PDB by Braxton and coworkers^[14]^ with PDB IDs 8G7J, 8G7L, and 8G7N, respectively, and in the **D_V72I-ADP-Pi_** case by replacing ATP with ADP-P_i_. Structures for **S_WT_**, **D_WT_**, **F_WT_** and **D_WT-ADP-Pi_** were straightforwardly reconstructed from their V72I equivalents by introducing point-mutations with the *PyMOL* modelling suite.^[21]^ Finally, **M_WT_** and **M_V72I_** were directly derived, respectively, from chain A of **S_WT_** and **S_V72I_**.

We opted not to model any extra residues other than those already present in each CryoEM structure: as such, all Hsp60 monomers in **D**, and **F** end with Glu526, whereas **M** and all **S** monomers end with Lys525. At the *N-*terminus, an extra alanine that precedes Ala1 in every monomer in all CryoEM structures and appears as “Ala0”^[14]^ (but is not actually part of mature Hsp60) was only deleted from **D**, and **F** but retained in **M** and **S**: this conveniently ensured that every monomer in every system retained a total of 526 residues. Similarly, all Hsp10 units in **F_WT_** / **F_V72I_** were modelled from residue Gly3 to residue Asp102, as present in the **F_V72I_** CryoEM structure (PDB ID: 8G7N).^[14]^

All Mg^2+^ and K^+^ cations present in the original CryoEM structures of the double ring and the football were retained in final **D_WT_** / **D_V72I_**, **D_WT-ADP-Pi_** / **D_V72I-ADP-Pi_** and **F_WT_** / **F_V72I_** structures.

Hydrogen atoms were added *via* the *reduce* utility in *AmberTools* (v. 21),^[22]^ which ruled out the presence of disulfide bridges in all systems, predicted the optimal orientation of Asn and Gln sidechains, and predicted His242 and His315 present in Hsp60 to be protonated on Nε2 in every monomer of every system (no histidines are present in Hsp10).

##### Protonation states of the catalytic Asp50/Asp397 dyad

Almost all titratable residues were modelled in their standard protonation state, *per* pK_a_ predictions by the *PropKa* package^[23]^ at physiological pH. A significant exception, however, was represented by the catalytically relevant^[24]^ Asp50 and Asp397 dyad. These aspartates are located, respectively, in the equatorial and intermediate domains of Hsp60 at either side of the ATP binding pocket (Figure 1b). In apo systems **M** and **S**, where pockets are open and solvent-exposed, both aspartates were predicted to be deprotonated, with low pK_a_ values (*e.g.*, 3.18 for Asp50 and 4.91 for Asp397 in **S_V72I_**). On the other hand, in **D** and **F** systems, where ATP is present with Mg^2+^ and K^+^, pockets are more closed and Asp50/Asp397 are closer together: here, predicted pK_a_ values increase significantly (*e.g.*, 8.22/12.52 in **D_V72I_** and 7.13/11.13 in **F_V72I_**).

Elevated pK_a_ values led us to anticipate that the two aspartates might act as a catalytic dyad in ATP hydrolysis, in which either Asp could be driven to act as the general base with the other Asp perhaps already protonated. It is for this reason that we expressly decided to consider structures of **D_WT_**, **D_V72I_**, **F_WT_**, and **F_V72I_** with Hsp60 monomers in one of three protonation states: (1) Asp50/Asp397 both deprotonated, in line with predictions for **M** and **S**; (2) Asp50 protonated; and (3) Asp397 protonated. Whenever we need to specifically refer to one of these protonation states, generic labels will be expanded to, *e.g.*, one of **D_WT-AspAsp_**, **D_WT-AshAsp_**, or **D_WT-AspAsh_**, as listed in Table 1. In addition, since we speculated Asp397 in the catalytic dyad could well become protonated as a result of ATP hydrolysis, we also chose to simulate two additional systems (labelled **D_WT-ADP-Pi_** / **D_V72I-ADP-Pi_**) in which all Asp397 residues in **D** are left protonated, but ATP is replaced by ADP and [HPO_4_]^2–^ (P_i_); (Mg^2+^ and K^+^ are left in place).

##### Capping and solvation

After construction and hydrogenation/protonation of our systems, *N-* and *C-*termini of each monomer were respectively capped with –NH_3_^+^ and –COO^−^ groups using the *tleap* utility in *AmberTools*.^3^ The tool was also employed to add likely crystallographic waters around the apical and equatorial domains of Hsp60, as determined by backbone-atom superimposition of two high-resolution crystal structures of bacterial GroEL homologues (respectively, PDB ID: 3VZ6 and PDB ID: 1KP8).^[25]^ Crystallographic waters around Hsp10 in **F** systems were similarly determined by superimposing bacterial GroES (PDB ID: 3NX6) as deposited by the Kang group.

Addition of crystallographic waters was followed by solvation in a cuboidal box of water molecules with edges at least 12 Å away from any **M**, **S**, **D**, or **F** atom. Finally, in order to neutralise excess negative charge, we again used *tleap*^3^ to randomly position an appropriate number of Na^+^ cations in the simulation box. Coordinates and topologies of all starting structures are provided as SI.

#### Atomistic Molecular Dynamics Simulations

##### Forcefield parameters

Standard protein residues including protonated aspartates are modelled using the *ff14SB* forcefield,^[26]^ whereas the TIP3P model is chosen for water molecules.^[27]^ Cations are modelled as follows: Na^+^ as prescribed by Joung and Cheatham (as well as K^+^ where present); ^[28]^ for Mg^2+^, where present, we instead employ parameters published by Allnér *et al.*^[29]^ ATP is modelled according to parameters published by Meagher and coworkers.^[30]^ In simulations of **D_ADP-Pi_**, ADP is treated according to parameters by the same authors,^[30]^ whereas forcefield parameters for [HPO_4_]^2–^ are those published by Kashef Ol Gheta and Vila Verde. ^[31]^

##### Main simulation details

Atomistic Molecular Dynamics (MD) simulations were run using the *AMBER* software (version 2020, www.ambermd.org), starting from each of the 18 structures listed in Table 1 and using a common recipe that comprised: preminimisation if necessary; minimisation; preproduction (2.069 ns); and production (500 ns). In fact, four independent MD replicas were run for each system—*i.e.*, with atomic velocities assigned each time from a different random seed prior to preproduction—for a total of 18 systems × 4 replicas = 72 total replicas, and a total production time of 500 ns × 4 replicas = 2 µs per system.

Preminimisation was carried out with the *ParmEd* tool^[32]^ on all **S**, **D**, and **F** variants, by applying 50-100 minimiser steps, depending on the system. Minimisation and early preproduction stages were carried out for all systems using the *sander* utility in *AMBER*; thereafter we switched to the *pmemd.cuda* utility, which benefits from GPU- acceleration.^[33]^

Full details regarding minimisation and preproduction are reported as SI, as are input scripts for each stage. The production stage itself integrates equations of motion with a timestep of 2 fs and is carried out in the *NpT* ensemble under periodic boundary conditions: the constant temperature of 300 K is enforced *via* Langevin’s thermostat (5.0 ps^−1^ collision frequency);^[34]^ the constant pressure is stabilised at 1.0 bar using Berendsen’s barostat^[35]^ (relaxation time 1.0 ps). The cutoff for the computation of Lennard-Jones (LJ) and Coulomb forces in direct space is set at 8 Å from each atom; beyond this cutoff, LJ interactions are truncated, whereas Coulomb interactions are calculated in reciprocal space. Coulomb interactions are evaluated according to the Particle Mesh Ewald approach.^[36]^ All positional restraints previously applied to backbone atoms (*cf.* SI) are lifted; note that K^+^ and Mg^2+^ cations present in Hsp60 units in **D** and **F** are never subjected to restraints except in the very first minimisation stage. On the other hand, all bonds containing hydrogen are constrained using the SHAKE approach.^[37]^

MD trajectory postprocessing and analysis, including concatenation into metatrajectories, key distance and radial distribution function (RDF) plots were performed using the *cpptraj* utility distributed with *AmberTools* (v. 21).^[22]^ 2D histograms were built and normalised from *cpptraj* data using the *seaborn* Python package.^[38]^

##### Distance Fluctuation Analysis (DF)

Distance Fluctuation (DF) analysis^[39]^ examines a MD (meta)trajectory with the objective of quantifying the degree of allosteric communication occurring between every residue pair in a protein/protein complex (“inter-residue crosstalk”), no matter how distant its constituent residues are from each other. This is useful as allosteric phenomena tend to occur over longer timescales than those typically accessible by unbiased MD. ^[39b, 40]^

The code to perform DF analysis is available online at https://github.com/colombolab/Distance-Fluctuation-DF-Analysis.

In this work, DF analysis is separately carried out on the full 2 µs MD metatrajectory of each system listed in Table 1 except **D_ADP-Pi_** (*i.e.*, the four concatenated 500 ns production stages of its MD replicas): the analysis only requires a metatrajectory of Cα atoms and requires no prior alignment or manipulation other than making sure that in **S**, **D**, and **F**, all individual monomers appear compactly within every MD frame (no periodic images). From the metatrajectory of a system with *N* residues, DF analysis yields a *N* × *N* **DF** matrix whose individual elements *DF_ij_* represent the degree of distance fluctuation (“DF score”) between the *i*^th^ and *j*^th^ residue. Each *DF_ij_* is derived as follows:

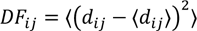

where *d_ij_* represents the distance between the two residues in a particular frame measured at their Cα atoms, and quantities comprised between ⟨⟩ are averages over the 40000 frames present in each metatrajectory (1 every 50 ps, 10000 per replica). Residue pairs with the lowest DF scores deviate the least from their average distance ⟨*d_ij_*⟩, and are associated with the highest degree of allosteric communication, moving as if connected through an imaginary rod; conversely, highest DF scores are associated with highly fluctuant residue pairs that exhibit poor allosteric communication.

Subtraction of DF matrices, one from a system with WT Hsp60 and the other from one with V72I, readily highlights regions gaining allosteric prominence upon mutation, and those losing it.

##### Shortest Path Map (SPM)

The shortest path map (SPM)^[41]^ eloquently charts the principal allosteric communication route(s) across a protein or multiprotein complex: in this work, we produce SPMs for **M_WT_** and **M_V72I_** starting from their respective MD metatrajectories. The process was handled by the *DynaComm.py* code.^[41]^

Broadly speaking, the first step entails the transformation of Hsp60 into a node-and-edge graph, with each residue becoming a node. An edge is drawn to connect nodes *i* and *j* (residues *i* and *j*) only if during the combined MD replicas Cα atoms of residues *i* and *j* remain, on average, closer than 6 Å. Furthermore, the length *l_ij_* of the edge is inversely proportional to the magnitude of (anti)correlation |*C_ij_*| between *i* and *j*, as shown by the formula

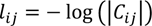

where correlated (*C_ij_* > 0), anticorrelated (*C_ij_* < 0), or uncorrelated movement (*C_ij_* = 0) between *i* and *j* is quantified by

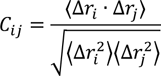

using the average displacements Δ*r_i_* and Δ*r_j_* of Cα atoms in *i* and *j* from their positions in the most populated conformational cluster. Thus, the shortest edges connect vicinal residues (≤6 Å) exhibiting the greatest correlation or anticorrelation.

The next step is to systematically trace the shortest path from each residue to every other residue prioritising travel across the shortest edges of the node-and-edge graph: in our case, we instructed the *DynaComm.py* code^24, 25^ to do this using the walktrap algorithm.^[42]^ As these shortest paths are being traced, the code cumulatively records how often individual edges are travelled through: at the end of the process, the most-often utilised edge is given a weight of 1, and the ensuing final SPM (projectable on the parent structure) is produced by only outputting those nodes that are connected by edges whose weight exceeds an arbitrary threshold of 0.3. To obtain SPMs from MD metatrajectories of **M_WT_** and **M_V72I_** all that was required beforehand was a pairwise (residue × residue) correlation matrix **C** (with elements *C_ij_*) calculated for each system using the most populated cluster as a reference; and an average distance matrix **d** (with elements ⟨*d_ij_*⟩). The (straightforward) derivation of **C** and **d** with *cpptraj*^17^ is reported as SI, along with the necessary scripts and the matrices themselves; the SPM output is also provided. The *DynaComm.py* code itself is now available online.^[43]^.

##### Principal Component Analysis (PCA)

A “reference” principal component analysis (PCA) of **M_WT_** was carried out on its production metatrajectory, using *cpptraj* to compute and diagonalise covariance matrices for Cα atom coordinates of residues Lys2 to Glu522, and to obtain movies of the first two resulting eigenvectors (*i.e.*, principal components). Indeed, we should recall that the first principal components (covariance eigenvectors) recapitulate the most dominant motions within a protein or protein complex.^[44]^

Subsequently, using this initial **M_WT_** PCA as a standard reference for Hsp60 motions, we turned to **M_V72I_** and to individual Hsp60 monomers in all remaining multimeric systems in Table 1 (except for **D_ADP-Pi_**): MD metatrajectories were systematically analysed and the motions of individual Hsp60 monomers were cumulatively projected onto the first 10 principal components of **M_WT_**. The choice to only consider Lys2:Cα– Glu522:Cα in every Hsp60 monomer examined allowed us to always focus on the same 521 atoms despite the different Hsp60 residue intervals modelled in the various systems (Table 1). Furthermore, both for our reference PCA and for subsequent projections, Cα atoms from the CryoEM structure of **M** (monomer A from PDB ID: 8G7J of an **S**)^1^ were employed as our universal alignment reference.

Input scripts to perform PCA for **M_WT_**, to make movies of its first 3 principal components, and to project **M_V72I_** on its first 10 principal components are provided as SI.

## Results and Discussion

### Hsp60 V72I mutation has a dramatic effect on chaperone structure and function(s)

The V72I mutation in the *hsp60* gene is linked to hereditary spastic paraplegia 13 (SPG13), an autosomal dominant disease characterised by spasticity and weakness in the lower extremities. This mutant has been shown to stabilise the Hsp60 structure and enhance steady-state ATPase activity^[14, 19]^. However, before beginning our computational studies to probe this mechanism, we first wanted to confirm the impact of V72I on structure and function (See Biochemical Methods in SI). Accordingly, we expressed and purified recombinant, human Hsp60 as the mature, mitochondrial form. For our biochemical characterisation, we compared three proteins: wild type Hsp60 (Hsp60^WT^), along with Hsp60^V72I^ and Hsp60^D3G^, a recessive mutant associated with a separate disease (MitCHAP-60)^[18a]^ By native polyacrylamide gel electrophoresis (native PAGE), Hsp60^WT^ was relatively unstable and sampled both the oligomeric and monomeric forms at a range of concentrations (Figure S1a). In contrast, Hsp60^V72I^ was a stable oligomer and Hsp60^D3G^ was primarily in the monomeric form (Figure S1a). Next, we compared the steady state ATPase activity of these mutations, showing that Hsp60^V72I^ has slightly elevated ATPase activity compared to Hsp60^WT^, while Hsp60^D3G^ has reduced activity (Figure S1b). Then, we titrated Hsp10 into these reactions to understand the effects of mutations on coupling to Hsp10. We found that Hsp10 elevated the ATPase activity of Hsp60^WT^, as expected, but that this activity was severely impaired for both mutants (Figure S1c). We noted that Hsp10 actually decreased ATPase activity of Hsp60^V72I^, likely because the protein is already an active, stable oligomer and binding to Hsp10 only slows cycling. Finally, we measured the ability of these proteins to refold denatured malate dehydrogenase (MDH), a native client of Hsp60. While Hsp60^WT^ was able to restore MDH activity (as measured by depletion of MDH’s substrate, NADH; see Biochemical Methods in SI), the Hsp60^D3G^ mutant was completely inactive and the Hsp60^V72I^ mutant was modestly impaired (Figure S1d). These results highlight the special case of Hsp60^V72I^, which has dramatic effects on structure, but is still a modestly active refolding machine, making it an ideal mutation for asking mechanistic questions by molecular dynamic simulations. Thus, these results, consistent with the literature,^[14, 19–20]^ show that the V72I mutation, despite being a rather modest change in a seemingly innocuous region of the protein, has a profound effect on oligomer stability, ATPase activity, and Hsp10 coupling.

In the subsections that follow, we report the analyses that we systematically performed on the full 2 µs MD metatrajectories for each of our 18 simulated systems (Table 1). Starting from variants of **M** and moving up to higher complexes, we focus on analysing how pervasively mutation V72I is able to alter intra- and inter-monomer allosteric communication (even between monomers that are distant from each other).

The shortest path map (SPM),^[41]^ distance fluctuation analysis (DF),^[39]^ and Principal component analysis (PCA)^[44]^ carried out on **M_WT_** and/or **M_V72I_** serve as a reference against which to monitor changes in movements and allostery in higher complexes. Wherever possible, ‘horizontal’ comparisons are also made across variants and protonation states within the same kind of complex, as these are of course specifically indicative of the effects brought about by the V72I mutation. Furthermore, given the crucial role played by ATP hydrolysis in driving the allosteric cycle and aggregation of Hsp60, we specifically focus on the impact V72I might have on catalytically relevant residues.

### M_WT_ and M_V72I_

#### Shortest Path Map

To analyse allosteric communication networks characterising **M_WT_** and **M_V72I_**, we first applied the Shortest Path Map (SPM) method by Osuna *et al.* ^[41]^ as described in the *Computational Methods* section. Resulting SPMs for **M_WT_** and **M_V72I_** are shown in Figure 2a. The SPM in **M_WT_** encompasses most of the important functional regions, revealing that they are allosterically interconnected: starting from Glu517 in the *C-*terminal α_18_–β_18_ loop of the equatorial domain (*cf.* Figure 1c), which is known to participate in inter-monomer interfaces in **S** (Figure 1a), it leads all the way to the bottom reaches of the apical domain. Along its path, the SPM spans important portions of equatorial domain helix α_18_, which is adjacent to mutation site Val72, as well as helix α_14_ in the intermediate domain, which is part of the substructure capping the ATP-site, i.e. the ATP lid (containing a catalytically crucial Asp397)^[7]^, and is itself involved in forming inter-Hsp60 interfaces in **S**, **D**, and **F** alike. Also spanned by the SPM on its way to the apical domain together with α_18_ are intermediate domain sheets β_5_ and β_13_. In short, all crucial centres of **M_WT_** are allosterically interconnected: from the interfaces that hold together **S_WT_** and higher complexes in the equatorial and intermediate domains, through catalytically relevant areas in the intermediate domain, and all the way up to the apical domain.

**Figure 2.**
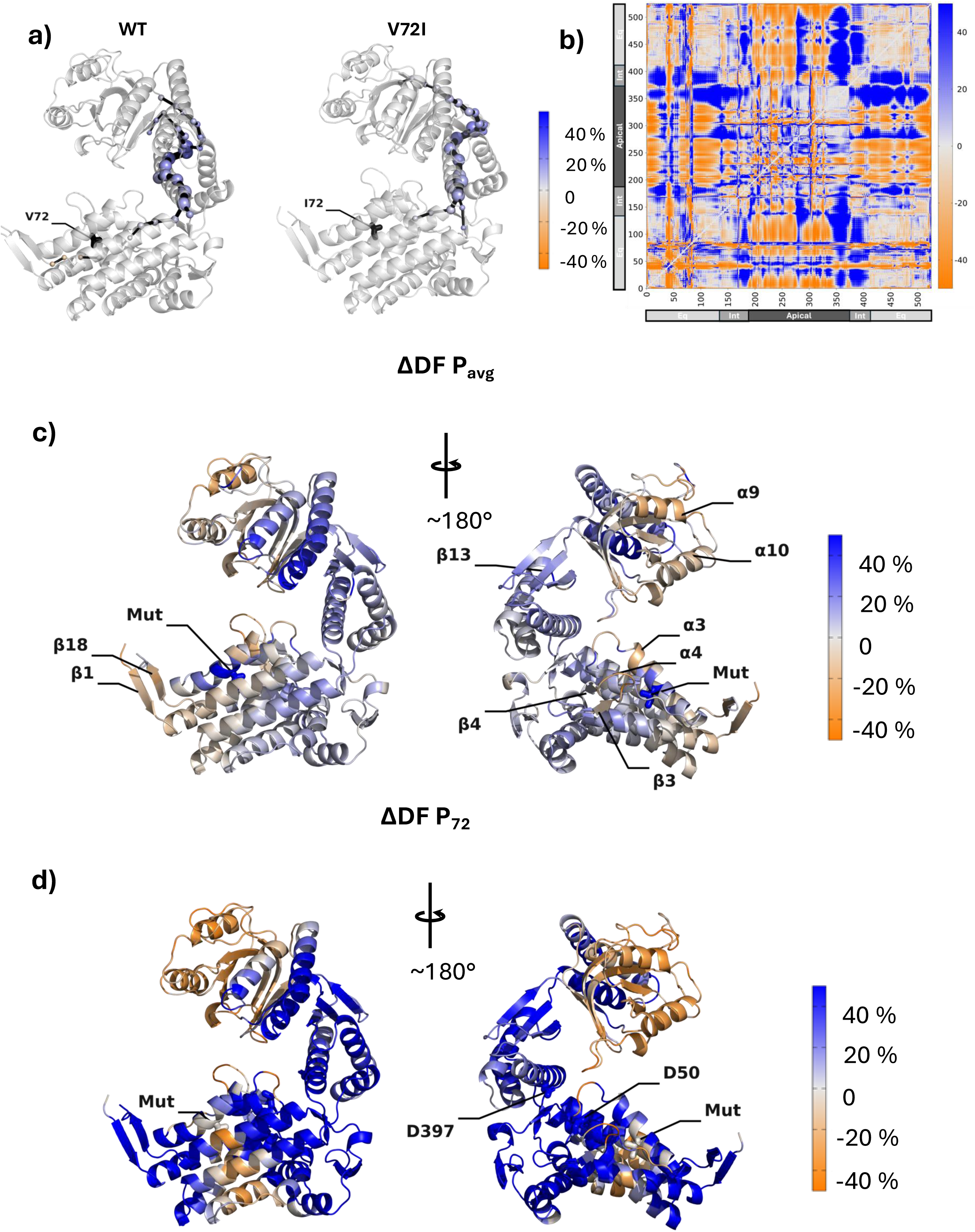
Distance Fluctuation (DF) and Shortest Path Map (SPM) analysis on. **M.** a) Shortest Path Map (SPM) of **M_WT_** and **M_V72I_**. Here SPM spheres are coloured according to the ΔDF P_avg_ value of the corresponding residues (see main text). The colour code is illustrated in the colour bar. b) Percentage DF difference matrix ΔDF obtained from DF_V72I_–DF_WT_ and taking DF_WT_ as reference for percentage calculations. Blue areas (positive values) correspond to loss of allosteric coordination in V72I whereas orange ones (negative values) indicate a gain in coordination in V72I. White areas are those unaffected by the mutation. Residue numbering is shown on the axis together with the corresponding domain: Equatorial (residues 0-133,410-525); Intermediate (residues 134-188,374-409); Apical (residues 189-373). To integrate this ΔDF matrix, as ESI we provide a video of each of its columns (one column = one residue) projected onto the starting structure of **M**. c) (P_avg_) Per-residue average change in DF score upon V72I mutation, projected onto the starting structure of **M**. The colour code is illustrated in the colour bar. d) (P_72_) Per-residue change in DF score with respect to residue 72, upon V72I mutation, projected onto the starting structure of **M**. The colour code is illustrated in the colour bar.

On the other hand, the SPM derived for **M_V72I_** featured in Figure 2a suggests that mutation of Val72 to Ile72 leads to a loss of ‘WT-like’ allosteric communication at crucial spots, particularly between the equatorial domain and the rest of the monomer. Specifically, the bulkier Ile72 sidechain significantly disrupts communication across helix α_18_ originating from the inter-monomer interface to the point that it almost entirely disappears from the SPM, were it not for a few of its *N-*terminal turns. Communication across the intermediate domain is similarly disrupted, as sheet β_13_ disappears from the path altogether and β_5_ alone is left to mediate communication between the equatorial and apical domains.

To recapitulate, therefore, comparison between the two SPMs clearly indicates that the extensive inter-domain allosteric communication that is present in **M_WT_** suffers an overall disruption in **M_V72I_**, becoming less efficient, and reducing communication across the intermediate domain.

#### DF Analysis

An alternative perspective on long-range effects in **M_WT_** and **M_V72I_** may be obtained by carrying out Distance Fluctuation analysis (DF)^[39]^ on residue Cα atoms, as described in the *Computational Methods* section. Pairwise DF matrices derived from MD metatrajectories of **M_WT_** and **M_V72I_** (Figure S2) will show low DF scores for residue pairs fluctuating in a highly coordinated manner, whereas residue pairs with high DF scores fluctuate more unrelatedly. Coordinated residue pairs, *i.e.*, those with low DF scores, are those giving the highest contribution in modulating functional motions; if residue pairs are distal, this indicates allosteric communication. As a reminder, if one subtracts the DF matrix derived from simulations of **M_WT_** from the one obtained from simulations of **M_V72I_** (*i.e.*, DF_V72I_–DF_WT_), the resulting difference matrix ΔDF (which can be normalised by taking percentages; Figure 2b) can eloquently provide insight into which residue pairs of **M** are allosterically disrupted by the mutation (positive values, lower coordination corresponding to less efficient allosteric message passing, blue areas in Figure 2b) and, conversely, which ones gain allosteric prominence because of it (negative values, higher coordination, orange areas). White areas are those unaffected by the mutation.

Normalising the ΔDF matrix in terms of percentage differences better highlights gains or losses of long-distance coordination, which is synonymous with allosteric prominence. Moreover, alongside the ΔDF matrix itself, it is particularly useful to examine two key projections of it onto the structure of **M_V72I_**, which we provide in Figures 2c and 2d (with each projection also shown rotated by 180°). Projection 1 (P_avg_) (Figure 2c) is that of the average change in DF score experienced by each residue upon mutation of Val72 to Ile (again, blue residues experience an average loss of allosteric coordination upon mutation, orange ones experience an overall gain). Projection 2 (P_72_) (Figure 2d), on the other hand, directly involves the 72^nd^ column of the ΔDF matrix, meaning that the colour of each residue (including Ile72 itself) specifically reflects whether that residue has gained (orange), lost (blue), or not changed (white) its level of allosteric coordination with Ile72 in **M_V72I_** compared to Val72. A video of all remaining projections P_0_ to P_525_ is provided as ESI for completeness.

Analysis of the percentage-normalised ΔDF matrix in Figure 2b and projections in Figures 2c-d reveal that the V72I mutation gives rise to some very interesting allosteric alterations that are not entirely captured by the SPM analysis (Figure 2a). To some extent, this is to be expected as the two methods are based on different approaches. Nonetheless, the general emerging picture is that of a general rewiring of the allosteric dialogue between domains, consistent with the literature.^[19–20, 45]^

Of particular interest is a generalised loss of allosteric coordination upon mutation, evidenced by the blue colouring, that is distinctly recognisable in P_avg_ and (especially) in P_72_ across most areas of the protein, and is particularly prominent in functionally relevant regions such as parts of the apical domain (with one notable exception), the ATP binding site, the ATP lid, and the equatorial β-sheets at Hsp60-Hsp60 interfaces (β_1_, β_2_, β_3_, β_18_). It is furthermore interesting to note that the entirety of residues constituting the ‘downregulated’ SPM in **M_V72I_** are seen to lose allosteric dialogue. This latter aspect, which certainly reinforces our DF findings, is easily verifiable since spheres along the two SPMs (*i.e.*, Cα atoms of constituent residues, Figure 2a) are given identical colours to those of the corresponding residues in P_avg_ (Figure 2c). Further to this point, again in agreement with SPM analysis, inspection of the ΔDF matrix (Figure 2b) reveals that one of the highest concentrations of blue (largest loss of dialogue) precisely involves that same intermediate domain β_13_ sheet that disappears from the SPM in **M_V72I_** (residues 375-380), especially in relation to the equatorial domain.

In contrast to this tendential disruption of allosteric communication pathways that are active in the WT chaperone, it is just as interesting to note that there are (orange) regions of **M** that consistently go in the opposite direction, undergoing notable gains in coordination upon mutation of Val72. First and foremost, this is true for helices α_9_ and α_10_ in the apical domain, both on average (P_avg_) and directly with respect to Ile72 (P_72_): these helices stand out because they are crucial in forming interfaces with Hsp10 in **F** and face towards the inner cavity of the Hsp60 folding chamber. They are also involved in forming Hsp60-Hsp60 interfaces, particularly in **D**. The other important region that gains allosteric prominence upon mutation are equatorial β-strands β_1_, β_2_ and β_18_. Crucially, these too are involved in the formation of the inter-Hsp60 interfaces and while this is only visible in P_avg_—meaning that the increase in allosteric coupling is only indirectly linked to Ile72—this finding is still very relevant, because it suggests that allosteric cross talk is not obliterated, but rather, it is shifted so that β_1_, β_2_ and β_18_ are now directly communicating with apical helices α_9_ and α_10_.

Indeed, from the extra (bidimensional) detail provided by the full ΔDF matrix (Figure 2b), it can be seen how the greatest gains in allosteric dialogue (*i.e.*, more intense shades of orange) for β_1_ (residues 2-7) and β_18_ (518-522) are precisely accumulated in α_9_ (231-240) and α_10_ (256-267) more than elsewhere, suggesting that mutant Hsp60 establishes a more direct allosteric communication route between equatorial interfacial β-sheets and apical interfacial helices α_9_–α_10_. Further supporting the hypothesised Ile72-driven allosteric reorganisation of **M** is the orange hue in P_avg_ (Figure 2c) of the equatorial α_3_-α_4_ loop which lies directly between interfacial equatorial β-strands and apical helices α_9_–α_10_. Another indication is that the β1-α1 residues are the only ‘orange’ residues in the SPM of **M_WT_**, and they disappear from the SPM of **M_V72I_** (Figure 2a; *vide supra*), indicating that their allosteric coordination plays a role only in the WT.

Finally, it is particularly eloquent to visualise P_avg_ and P_72_ with reference to position 72. In the former case (Figure 2c), Ile72 is the bluest residue of all, signalling that it has a much diminished level of allosteric dialogue, on average, compared to Val72. In P_72_ (Figure 2d), the only notably ‘orange’ area outside the apical domain is a rough ‘sphere’ surrounding Ile72, suggesting a strong allosteric perturbation (local gain) of the Ile sidechain on vicinal residues. It is clear, again, how most other areas lose their allosteric connection to Ile72.

In summary, consistently with SPM analysis, DF analysis confirms that the point mutation of Val72 to Ile72 has pervasive effects on the internal, functionally oriented dynamics of the Hsp60 monomer, whose mechanisms of allosteric coordination appear to be significantly rewired/rerouted. Compared to **M_WT_**, we observe a significant loss in allosteric cross-talk between the equatorial domain, the catalytic ATP lid/intermediate domain, and non-interfacial regions of the apical domain. In parallel, there is evidence of the emergence of a new set of distinct coordination patterns, in which equatorial interfacial β-sheets directly dialogue with the interfacial helices α_9_ and α_10_, involved in the assembly of Hsp60-Hsp60, Hsp60-Hsp10, and Hsp60-client interfaces. This mechanism may favour the preorganisation of the functional interfacial substructures for the recognition of other partners in the formation of the larger functional assemblies. We also note that what we observe here is in line with the rigidification of mutant Hsp60 that is observed experimentally.^[14, 19, 45]^

We will look for further clues to this allosteric rewiring in the analyses that follow for **M** and higher complexes.

#### Principal component analysis and conformational descriptors

To further characterise differences in motion between **M_WT_** and **M_V72I_**, we first performed Principal Component Analysis (PCA) on the MD trajectory of **M_WT_**, taking it as our reference for Hsp60 monomer dynamics (see *Computational Methods*). Only the first two principal components, henceforth PC1 and PC2, were taken into account for subsequent analysis. PC1 describes opening-closing of the apical domain, while PC2 captures the counterclockwise rotation of the apical domain with respect to the intermediate domain β-sheets, together with α_14_ ATP lid closure movements. Movies of both PC1 and PC2 (from the most negative values to the most positive) are provided as ESI.

Projection of the metatrajectories of **M_WT_** and **M_V72I_** onto these two principal components of **M_WT_** (Figure 3a) shows important differences between the two, with the mutant sampling a more confined region in a way that is in line with experimentally predicted rigidification, and consistent with a rewired allostery. The greater freedom of movement of **M_WT_** is particularly evident at values of PC1 +25 to +100 (apical opening): while a substantial minority of **M_WT_** simulation frames manage to sample this area, simulations of **M_V72I_** reaching it are clearly scarcer. Furthermore, the subset of more open conformations simulated for **M_WT_** distinctly envelops the square dot, which represents monomers in double ring **D**, whereas **M_V72I_** simulations leave this region unexplored. In short, it is easier for **M_WT_** to directly adopt ATP binding conformations seen in **D** than it is for **M_V72I_**. Along PC2, **M_V72I_** alone is able to sample values of PC2 below –50, indicating that apical domains are less prone to rotate about the intermediate domains: this is again consistent with attenuation of allosteric pathways across the intermediate domain.

**Figure 3.**
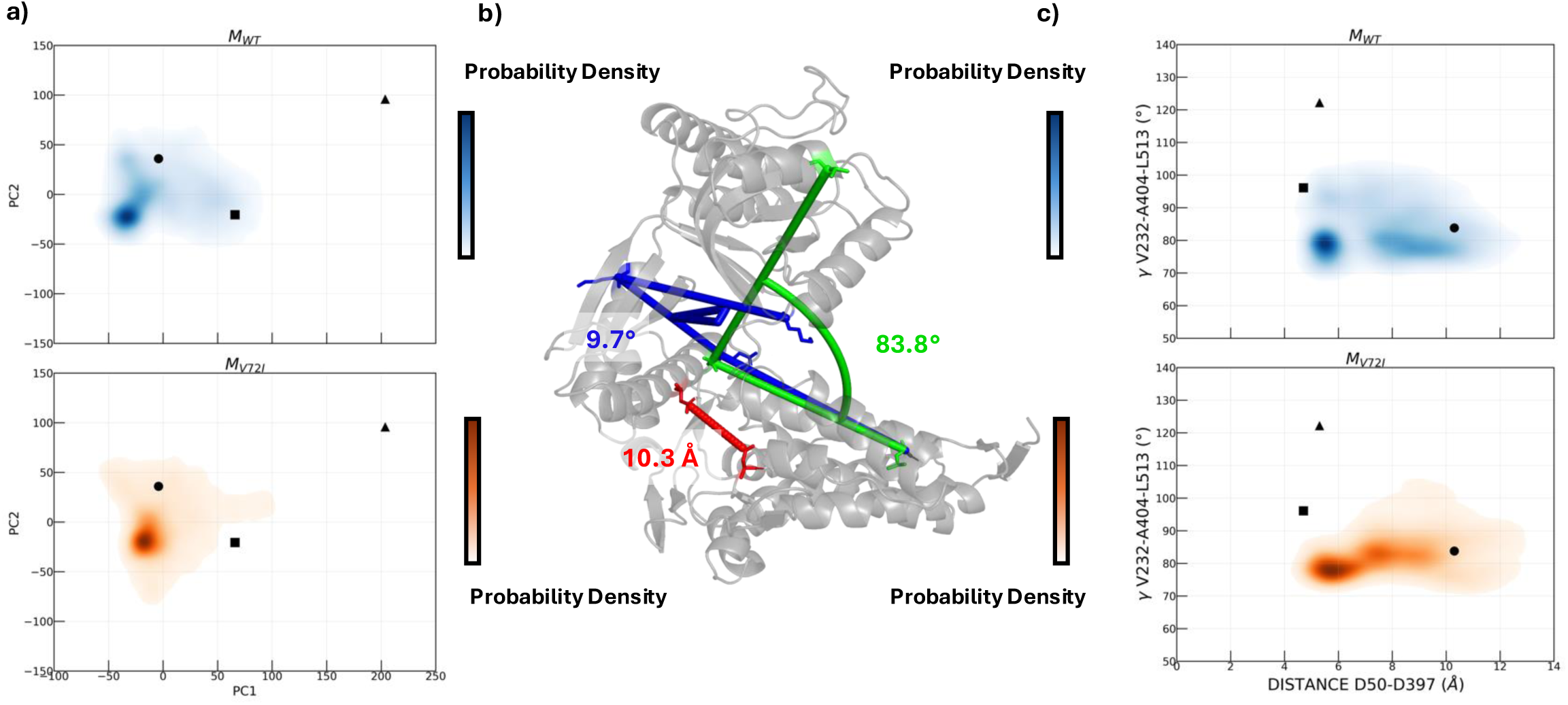
Motions in M during MD simulations. a) Projection of MD metatrajectories of **M_WT_** (top; blue) and **M_V72I_** (bottom; orange) onto principal components PC1 and PC2 as derived from MD metatrajectories of **M_WT_** (see main text). b) Structural variables best describing the main collective motions and reactivity of **M**, marked on its starting structure with their respective values (atoms defining these are reported in the main text). Green: angle **γ** (apical domain opening in PC1); blue: dihedral **χ** (apical domain counterclockwise rotation in PC2); red: distance **δ** between catalytically relevant aspartates. c) Distribution of **δ** and **γ** across MD metatrajectories of **M_WT_** (top; blue) and **M_V72I_** (bottom; orange). In panels a) and c), round dots represent values of plotted quantities in the CryoEM structure of **S** (**M**); square dots values in the structure of **D**; triangular dots values in the structure of **F**. Colour intensity is proportional to the probability density function, which is directly proportional to the number of MD frames with a given set of values.

As a general conformational trait of **M**, it is finally interesting to note the two distributions with respect to the round dot in Figure 3a, which represents PC1 and PC2 of all monomers in **M_WT_** and **M_V72I_**’s parent CryoEM structure of **S_V72I_**. The fact that the bulk of **M_WT_** and **M_V72I_** frames concentrates away from the dot, at lower PC1 and PC2 values, indicates that as soon as **M** is isolated from **S**, it tends to ‘close up’ significantly.

The observation of such clear-cut principal motions prompted us to identify simpler structural variables to describe conformational mechanisms in the more complex assemblies discussed in the following paragraphs. These are illustrated for **M** in Figure 3b.

Specifically, we selected the angle between Cα atoms of residues Val232, Ala404 and Leu513 (**γ**), which recapitulates the opening of the angle between the two domains captured by PC1, and the dihedral defined by Cα atoms of Ala8, Thr28, Glu255 and Lys266 (**χ**), which is used to describe the rotation evidenced by PC2. To these, we also added the distance (**δ**) between Cγ atoms of Asp50 and Asp397 forming the catalytic dyad, to account for structural modifications that may impact ATP processing. In particular, distribution plots of the angle **γ** and the distance **δ** along the MD metatrajectories of **M_WT_** and **M_V72I_** highlighted the most significant variations in the dynamics of the two proteins (Figure 3c). Interestingly, just like PC1/PC2 plots in Figure 3a, these plots reveal that only **M_WT_** just about populates values of **δ** and **γ** including those observed for the Cryo-EM structure of the ATP-bound double ring **D** (square dot). This does not happen for **M_V72I_**, which is clearly less favoured to explore **γ** values above 90°, in accordance with the more negative values of PC1. It is also evident that **M_V72I_** favours lower values of **δ**: both **M_WT_** and **M_V72I_** feature a main peak centred at around 5.75 Å, but in the latter case, there are instances in which Asp50:Cγ and Asp397:Cγ are able to come as close as 4.25 Å. This may be an early indication of a more suitable positioning of the catalytic residues for reactivity. Overall, these findings again point to a rigidification of **M_V72I_** dynamics, in line with our previous observations and with experimental evidence.

Identified descriptors **δ** and **γ** together with the projections of Cα metatrajectories of the various complexes onto the space defined by **M_WT_** PC1 and PC2 will be instrumental in the following subsections to describe and compare the dynamics of **S**, **D** and **F** and characterise possible effects of the V72I mutation.

#### S_WT_ and S_V72I_

We first set out to analyse the dynamics of the WT and V72I variants of Hsp60 when assembled in apo 7-meric single ring complexes **S_WT_** and **S_V72I_** (Figure 1a). Specifically, we first qualitatively characterise the main conformational variations of individual monomers with respect to **M_WT_** (PCA), we then move to the analysis of the critical distance between the catalytic aspartates, and finally, we investigate the overall internal dynamics via DF analysis.

#### Principal component analysis and conformational descriptors

Compared to isolated **M**, PCA shows that individual **S** monomers explore a smaller but denser interval along PC1, hardly ever exceeding 0 (Figure 4a). This is consistently reflected by apical domains that remain tendentially closed in both the WT and V72I variants, as testified by values of **γ** in Figure 4b that are locked below 90°, and shows that apical domains are not able to open in **S**. Again in both variants (Figure 4b), **δ** is uniformly distributed around 10 Å, signalling that the arrangement of catalytic aspartates in **S** remains unreactive, as expected in the absence of ATP.

**Figure 4.**
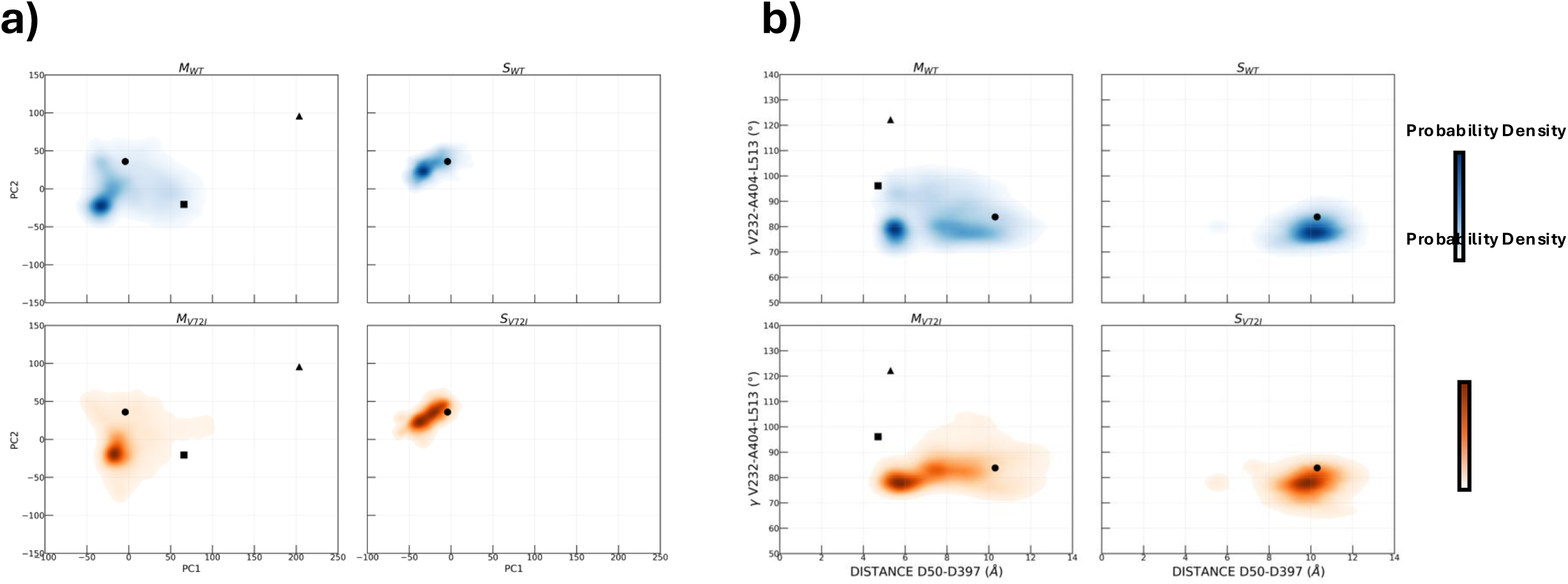
Motions in S during MD simulations. a) Projection of MD metatrajectories of **S_WT_** (cumulative 7-monomer projections; top right; blue), and **S_V72I_** (cumulative 7-monomer projections; bottom right; orange) onto principal components PC1 and PC2 as derived from MD metatrajectories of **M_WT_** (see main text). Projections of MD metatrajectories of **M_WT_** (top left; blue), and **M_V72I_** (bottom left; orange) are retained for comparison. Round dots represent values of PC1 and PC2 in the CryoEM structure of **S** (**M**); square dots (left only) values in the structure of **D**; triangular dots (left only) values in the structure of **F**. b) Distribution of distance **δ** and angle **γ** (see main text) in MD metatrajectories of **S_WT_** (cumulative counts for individual monomers; top right; blue), and **S_V72I_** (cumulative counts for individual monomers; bottom right; orange). Distribution in **M_WT_** (top left; blue), and **M_V72I_** (bottom left; orange) is retained for comparison. Round dots represent values of **δ** and **γ** in the CryoEM structure of **S** (**M**); square dots (left only) values in the structure of **D**; triangular dots (left only) values in the structure of **F**. Colour intensity in both panels is proportional to the probability density function, which is directly proportional to the number of MD frames with a given set of values.

The heptameric arrangement thus appears to restrain the conformational freedom of individual monomers compared to the unassembled state, favouring a rigid-open state, and an open ATP pocket. This is indistinctly true for both WT and V72I Hsp60.

#### DF analysis

Distance Fluctuation analysis (DF) was carried out on metatrajectories of **S_WT_** and **S_V72I_** following the same conceptual scheme as described in the section dedicated to **M**, and in this case, some allosteric differences between **S_WT_** and **S_V72I_** begin to emerge. P_avg_ (Figure 5a) shows that in this case too, mutation of Val72 to Ile generally tends to rewire allosteric coordination, which results in a complex spectrum: this is to be expected for a large system whose multifaceted dynamics aptly reflects its multimeric composition.

**Figure 5.**
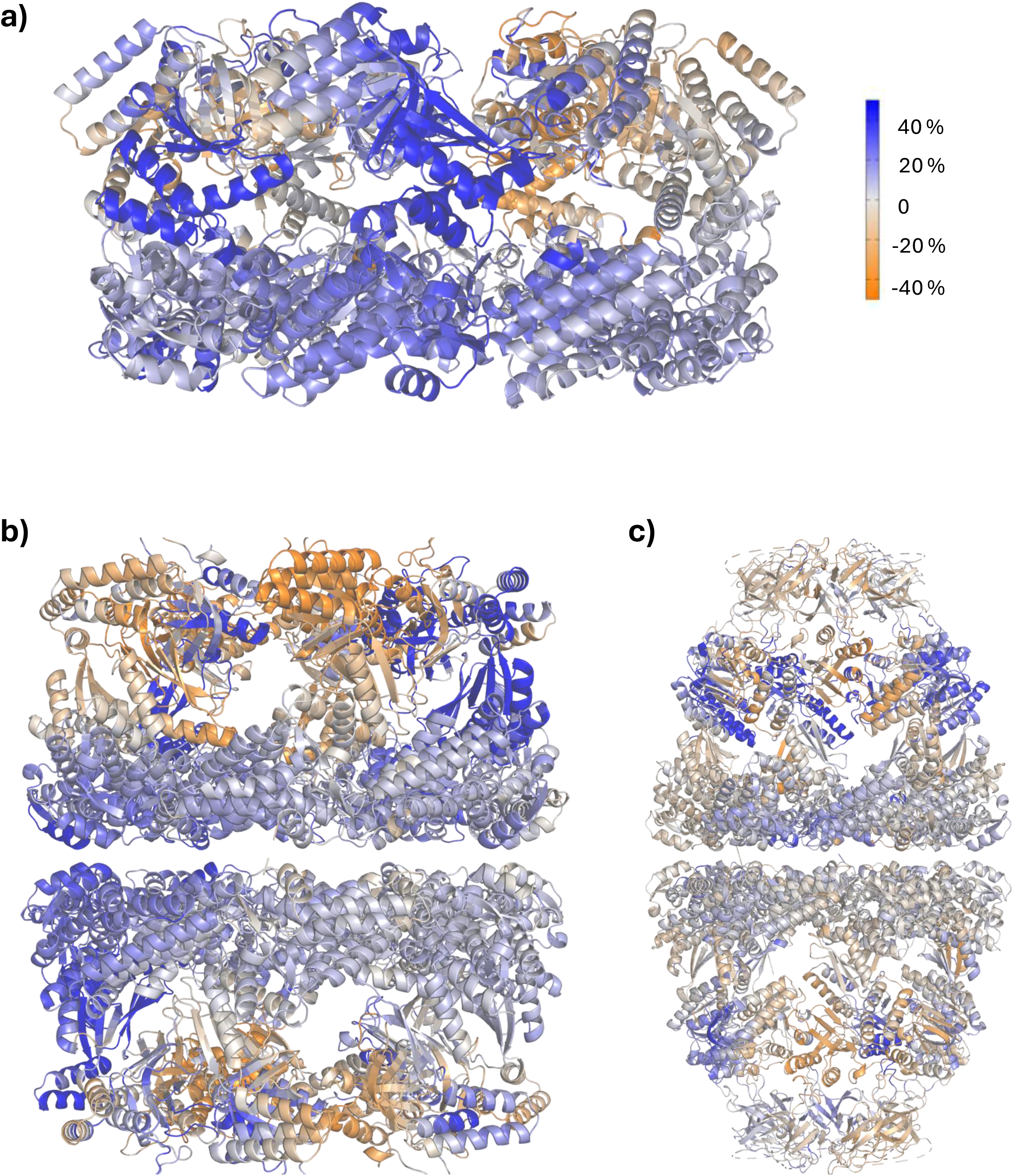
Distance Fluctuation (DF) analysis on S, D, and F. a) Average percentage change in DF score for each residue upon V72I mutation projected onto the starting structure of **S** (P_avg_). b) Average percentage change in DF score for each residue upon V72I mutation projected onto the starting structure of **D** (P_avg_). The change is calculated from simulations of **D_WT-AspAsh_** and **D_V72I-AspAsh_**. c) Average percentage change in DF score for each residue upon V72I mutation projected onto the starting structure of **F** (P_avg_). The change is calculated from simulations of **F_WT-AspAsh_** and **F_V72I-AspAsh_**.

A glance at Figure 5a reveals a general prevalence of blue areas (coordination loss upon mutation) across equatorial and—especially—intermediate domains: this is in places qualitatively reminiscent of P_avg_ and P_72_ for **M** (Figures 2c-d) and confirms disruption of allosteric communication across the intermediate domains. Even more interestingly, there emerges an asymmetric alternation in the dynamic coordination of the apical domains, whereby in some monomers a preponderance of blue (loss of allosteric coordination) is observed, while in others a prevalent gain of allosteric coordination appears (orange in Figure 5a). The results of DF analysis are particularly interesting as this picture reflects the asymmetric conformational patterns observed by Braxton and coworkers:^[14]^ the tendency for some monomers in the complex (the ones with lower allosteric coordination) to explore alternative conformations may in fact favour their upward motions to form contacts with the cochaperones carrying the client.

We reiterate that, in general, P_avg_ for **S_WT_** and **S_V72_** qualitatively replicates what was observed in the monomers. It is worth noting here, however, that it is not possible to directly compare the results for the monomers in **S** with what observed for isolated **M**, given the higher complexity of the system. The presence of contacts with the other constituent monomers significantly affects the specific intramolecular degrees of coordination of residue 72 with the rest of the (whole) system. For this reason, individual P_72_s will not be examined.

#### D_WT_ and D_V72I_

Double rings **D** are formed by two single rings **S** that are equatorially interfaced (Figure 1a), with the crucial inclusion of one ATP molecule, one Mg^2+^, and one K^+^ in each of the 14 active sites; these are all resolved in the experimentally characterised structures. Arrival of cations and ATP triggers an opening and rotation of the apical domain, and the closure of intermediate domain helix α_14_ to lock catalytic Asp397 into place (Figure 1a); this change is also appreciable from the distinctive positions of the square dot in Figures 3-4 and some of the later Figures.

#### Principal Component Analysis

First, to understand the impact of V72I (and its extent) on the dynamics of **D_WT_** and **D_V72I_**, we set out to characterise them following the same scheme as shown in previous sections. Qualitatively, PCA analyses (Figure 6a) show that the overall essential space spanned by the monomers in both variants—and regardless of protonation state—is generally similar to the one spanned by the isolated monomers, especially along PC2. PC1 spans the same ranges as isolated **M_WT_** and **M_V72I_**, but both **D_WT_** and **D_V72I_** tend to sample more open conformations, as testified by the greater statistical weight of PC1 values above 0. This is, expectedly, in line with the more open starting structure (*cf.* square dots in Figure 6a). Further on the issue of apical openness in **D**, however, an important ‘horizontal’ difference (*i.e.*, between WT and V72I variants) does emerge with values of 0 ≤ PC1 ≤ +100 in two of the three protonation states examined, namely Asp50-Asp397 and Asp50-Ash397 (Figure 6a; second and fourth panels from the right). Monomers in mutant **D_V72I-AspAsp_** and **D_V72I-AspAsh_** tend to remain more ‘locked’ in their open starting conformations, with peaks along PC1 concentrating closer to the square dot representing the CryoEM of **D**. Conversely, wild-type monomers in **D_WT- AspAsp_** and **D_WT-AspAsh_** tend to move towards more closed apical domains (lower PC1 values) and appear to be in general more flexible.

**Figure 6.**
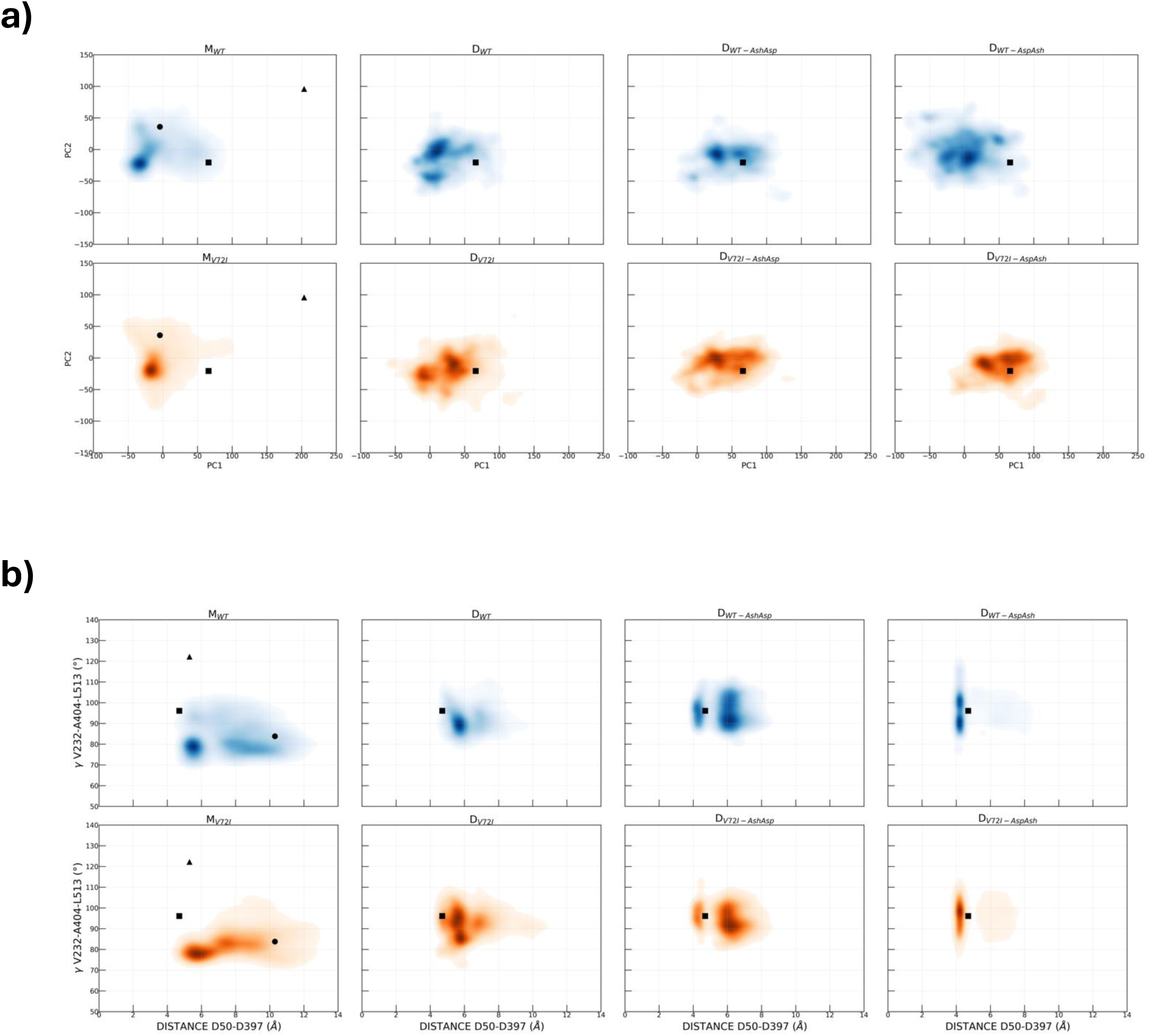
Motions in D during MD simulations. a) Projection of MD metatrajectories of **D_WT_** in three protonation states (cumulative 14-monomer projections; top 3 rightmost panels; blue), and **D_V72I_** in three protonation states (cumulative 14-monomer projections; bottom 3 rightmost panels; orange) onto principal components PC1 and PC2 as derived from MD metatrajectories of **M_WT_** (see main text). Second panels from the left pertain to simulations of **D_AspAsp_** (see main text); third panels from the left to simulations of **D_AshAsp_**; rightmost panels to simulations of **D_AspAsh_**. Projections of MD metatrajectories of **M_WT_** (top leftmost panel; blue), and **M_V72I_** (bottom leftmost panel; orange), are retained for comparison. b) Distribution of distance **δ** and angle **γ** (see main text) in MD metatrajectories of **D_WT_** in three protonation states (cumulative counts for individual monomers; top 3 rightmost panels; blue), and **D_V72I_** in three protonation states (cumulative counts for individual monomers; bottom 3 rightmost panels; orange). Second panels from the left pertain to simulations of **D_AspAsp_** (see main text); third panels from the left to simulations of **D_AshAsp_**; rightmost panels to simulations of **D_AspAsh_**. Distribution in **M_WT_** (top left; blue), and **M_V72I_** (bottom left; orange) is retained for comparison. In all panels, square, and triangular dots represent CryoEM structure values as described in Figures 3-4. As in Figures 3-4, colour intensity is proportional to the probability density function, which is directly proportional to the number of MD frames with a given set of values.

#### Conformational descriptor γ

The above difference is confirmed in the corresponding analysis of **δ** *vs.* **γ** that is plotted in Figure 6b. Focussing on **γ**, which we recall is closely associated with PC1 apical opening, it is clear from the rightmost panels that while **D_V72I-AspAsh_** appears to be significantly populating the ensemble around **γ** = 100°, **D_WT-AspAsh_** features an additional peak at **γ** = 90°, signalling some degree of closure and a second ‘preferred conformation’. Similarly, both fully deprotonated variants **D_WT-AspAsp_** and **D_V72I-AspAsp_** feature a prominent peak in the **γ** = 85° to 90° range, but **D_V72I-AspAsp_** additionally presents a second peak at around **γ** = 95°, signalling its proneness to remain more open. The possibility that WT has to transition to different conformations with lower barriers could on the one hand decrease the stability of the assembly, as testified by the greater difficulty in obtaining stable CryoEM structures. At the same time, it could favour its conformational adaptability to cochaperone Hsp10 and substrate proteins, an observation that may help rationalise the better ability of the WT protein to fold clients compared to the mutant. All of this is in line with what reported experimentally.^[14]^

Regardless of the variant, in any case, it is unsurprisingly confirmed by these plots too that **D** monomers have generally wider **γ** angles compared to isolated **M**. In this context, it is even more interesting to observe that in **D** Hsp60 monomers appear to populate different ensembles, with apical domains alternating in an up or down conformation. This result is again reminiscent of what observed by Braxton and co-workers^[14]^ and may indicate a tendency for the supramolecular complex to preorganise to recognise the client, which CryoEM structures show in contact with the down-conformations (Figure 7a-b).

**Figure 7.**
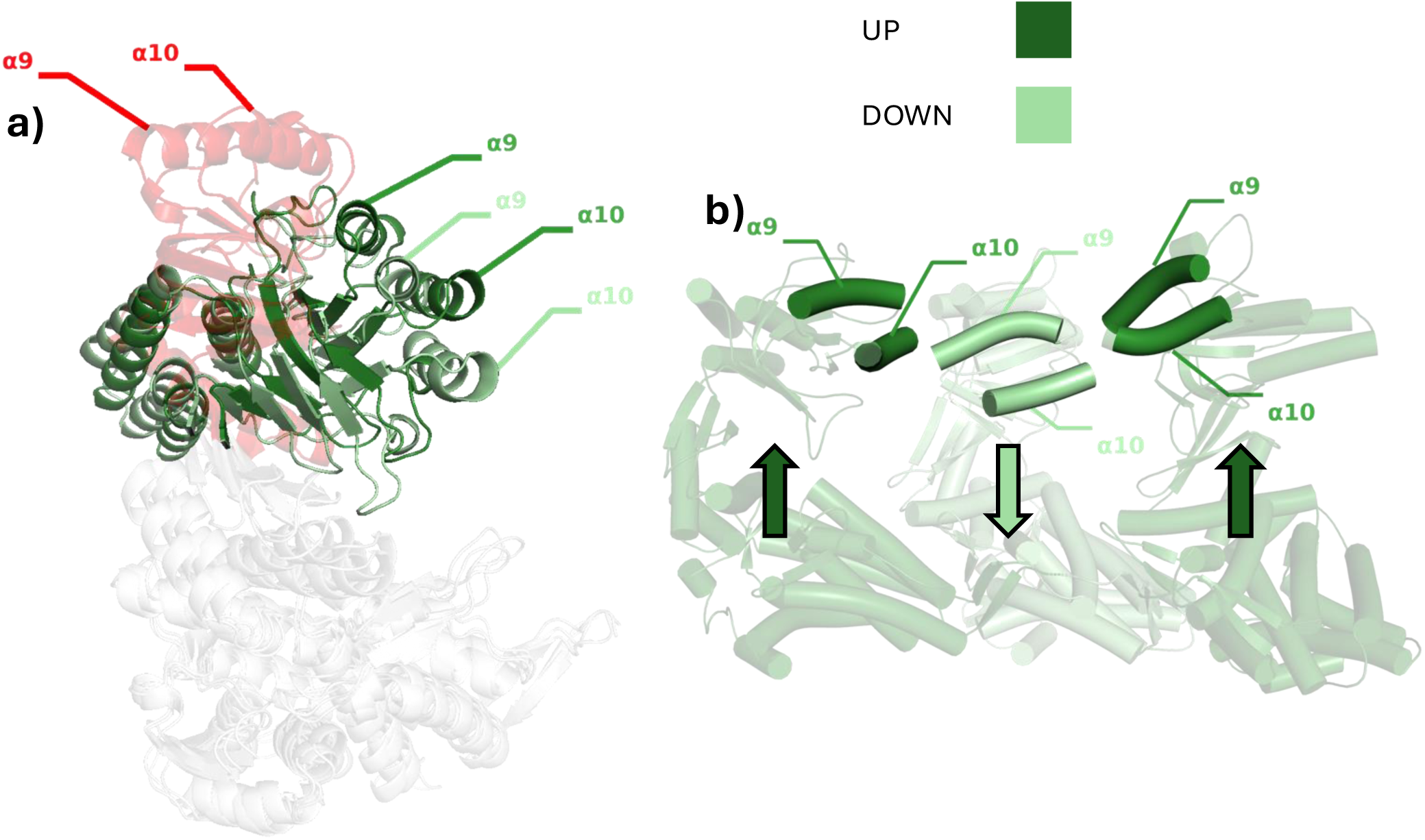
Up *vs.* down configurations of apical domains. a) Structures of 𝛼-helices 9 and 10 of the Hsp60 monomers involved in the up-down movement. Two **D_WT-AspAsh_** frames from MD are represented in green (lighter green down and darker green up), and **F_WT_** in red. Equatorial and intermediate domains are coloured in white. 𝛼-helix labels are depicted using the same colour of the reference structures. The “up” **D_WT-AspAsh_** pose has an angle of 103.7°, while the “down” **D_WT-AspAsh_** frame is 87.4° (the angle in **F_WT_** cryoEM structure is 122.2°). b) Structure of 3 adjacent monomers from a frame of MD of **D_WT-AspAsh_** (lower ring). As in the a panel, the “down” monomer is depicted in lighter green and “up” monomers in darker green (see the legend above). Again, 𝛼-helices 9 and 10 involved in the interactions with Hsp10 are highlighted, while the other domains of the monomers are transparent.

#### Conformational descriptor δ and reactivity

Another aspect that is conveniently assessable from simulations of **D** is any differences in the behaviour of the catalytic dyad around the natural substrate (with 14-fold statistics), and any influence on reactivity not only by the V72I mutation but also by the protonation patterns of the catalytic aspartates. Indeed, for this complex and for **F** specifically, the reader will recall that we noticed unusually high pK_a_ values for Asp50 and Asp397 (see *Computational Methods*) and this naturally prompted us to explore other protonation states of the catalytic dyad other than double deprotonation: either with Asp50 protonated, or with Asp397 protonated (Table 1).

To look more closely at the role of protonation *vs.* mutation in **D_WT_** and **D_V72I_**, we return to plots in Figure 6b, focussing this time on the Asp50:Cγ-Asp397:Cγ distance **δ**; Figure 6b plots are integrated with radial distribution functions (RDFs) of **δ** presented as ESI (Figures S3-S5). When both aspartates are deprotonated (third panel from the right and Figure S3), the total –2 charge pushes aspartate Cγ atoms apart to around 5.5 Å: importantly, however, in **D_V72I-AspAsp_** there emerges a distinctive “shoulder” at about 4.75 Å which—*cf.* square dot in the relevant Figure 8 panel—precisely corresponds to the (mutant) CryoEM distance. This signals that the mutation improves the chances of **D_V72I-AspAsp_** monomers to maintain the catalytic dyad closer together than **D_WT-AspAsp_**, and therefore suggests that in the absence of Hsp10, reactivity is slightly more enhanced in mutant Hsp60.

**Figure 8.**
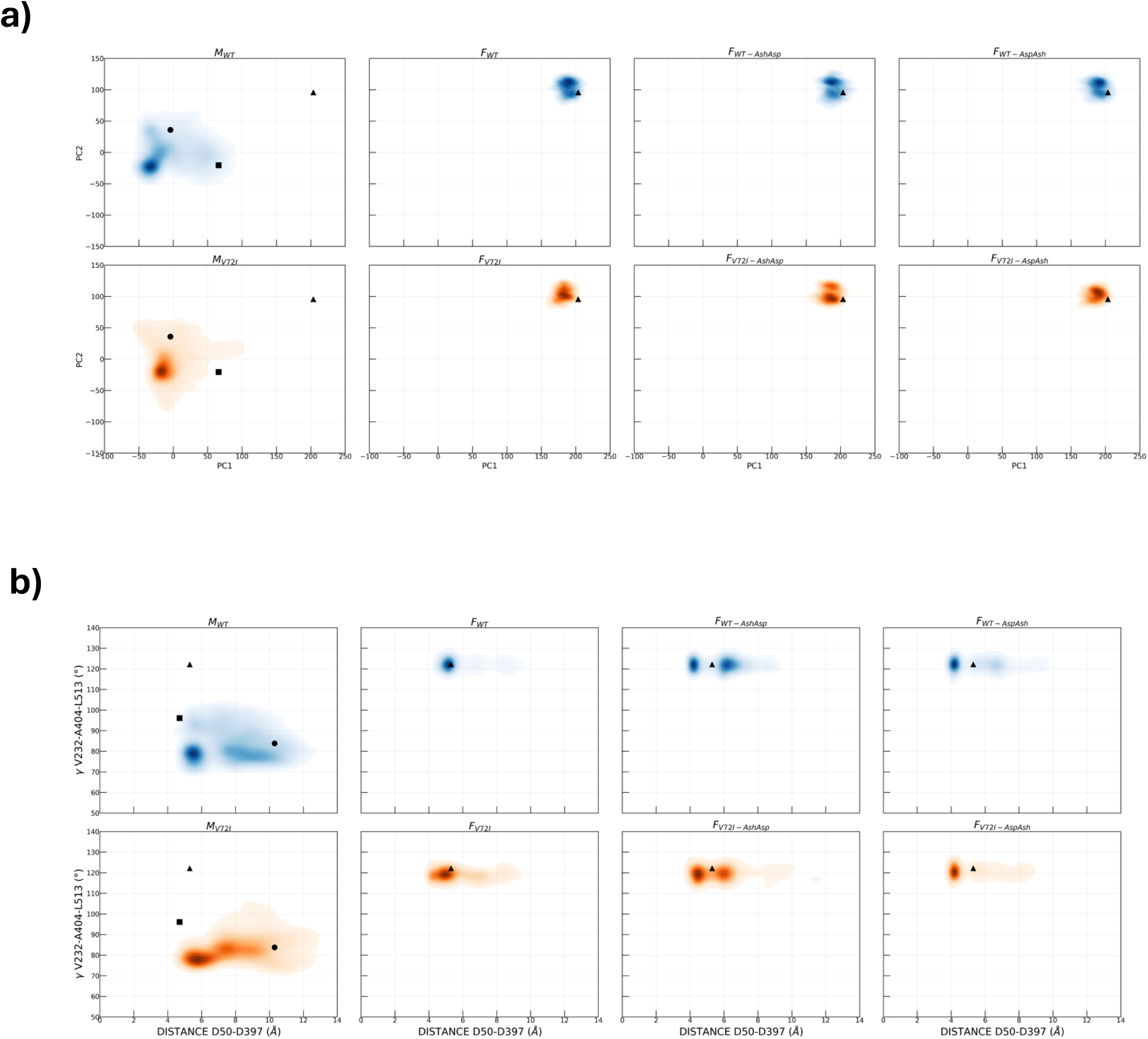
Motions in F during MD simulations. a) Projection of MD metatrajectories of **F_WT_** in three protonation states (cumulative projections for individual Hsp60 monomers; top 3 rightmost panels; blue), and **F_V72I_** in three protonation states (cumulative projections for individual Hsp60 monomers; bottom 3 rightmost panels; orange) onto principal components PC1 and PC2 as derived from MD metatrajectories of **M_WT_** (see main text). Second panels from the left pertain to simulations of **F_AspAsp_** (see main text); third panels from the left to simulations of **F_AshAsp_**; rightmost panels to simulations of **F_AspAsh_**. Projections of MD metatrajectories of **M_WT_** (top leftmost panel; blue), and **M_V72I_** (bottom leftmost panel; orange), are retained for comparison. b) Distribution of distance **δ** and angle **γ** (see main text) in MD metatrajectories of **F_WT_** in three protonation states (cumulative counts for individual Hsp60 monomers; top 3 rightmost panels; blue), and **F_V72I_** in three protonation states (cumulative counts for individual monomers; bottom 3 rightmost panels; orange). Second panels from the left pertain to simulations of **F_AspAsp_** (see main text); third panels from the left to simulations of **F_AshAsp_**; rightmost panels to simulations of **F_AspAsh_**. Distribution in **M_WT_** (top left; blue), and **M_V72I_** (bottom left; orange) is retained for comparison. In all panels, round, square, and triangular dots represent CryoEM structure values as described in Figures 3,4,6. As in Figures 3,4,6, colour intensity is proportional to the probability density function, which is directly proportional to the number of MD frames with a given set of values.

When a proton is introduced, **δ** plots and corresponding RDFs indicate that in these cases aspartates come closer. This is to be expected since the total charge is lowered to –1, and a hydrogen bond can be formed by holding aspartates together. Rather, it is more interesting to examine the important differences that are retained in the distributions of **δ** in **D_WT_** *vs.* **D_V72I_**. Upon protonation of Asp397 (rightmost panel and Figure S5), a sharp peak appears around 4 Å and a peak that was present at around 7 Å all but disappears. In the mutant, as is best seen from Figure S5), we note again that there is a higher frequency of poses in which aspartates are kept closer (integral difference ∼0.02). When protonation is introduced on Asp50 (second panel from right and Figure S4), the peak at 4 Å again appears, but there remains a second peak at 6 Å, and this time aspartates are kept closer in the WT variant.

In the closing subsection, we will better contextualise and discuss these important WT-V72I differences across protonation states, and discuss potential implications on reactivity. All we note at this stage is that they appear to be consistent from WT to V72I, and at the same time unrelated to differences in **γ**.

#### DF analysis

Here, we compare Distance Fluctuation analysis for simulations of **D_WT_** and **D_V72I_** across protonation states, to assess how mutation and protonation affect functional dynamics and allosteric coordination within the entire complex. Focussing mainly on the P_avg_ of **D_AspAsp_** (Figure S6) and **D_AspAsh_** (Figure 5b), one can recognise a marked increase in long range coordination (orange) concentrating across certain apical domains, particularly in the latter case, when Asp397 is protonated. In other words, the mutation consistently increases the allosteric coupling of some (not all) apical domains. Moreover, alongside the alternation of coordination in the apical domains and consistent with what observed in **S** and **M**, it is possible to appreciate the loss of coordination in the equatorial domains and almost all intermediate domains. Again, these specific dynamic alterations are expected to have repercussions on the ability of mutant **D** to interact with cochaperones and activate clients. As in **S**, individual P_72_s are not assessed.

To summarise, our data show that while **D_WT_** and **D_V72I_** share functional conformational states required to recruit the client and Hsp10, and to initiate ATP hydrolysis, the mutation is definitely observed to fine-tune all of these aspects across the complex. Its effects thus extend far beyond individual monomers. Mutant-induced modification of protein dynamics reverberates in the ability of assembled large complexes to interact with partners, contributing to the emergence of a modified functional profile. This expectedly translates into a cooperative modification of populations of the domains presented for interactions in large complexes and in this mode it is easy to see how a single mutation may significantly impact the organisation of the interaction networks that define cellular phenotypes, favouring pathologic traits.

#### F_WT_ and F_V72I_

Football-shaped complexes, **F**, represent the largest complex formed by Hsp60 studied in this work. This structure (Figure 1a) is formed once **D** further lifts its apical domains and recruits 7 Hsp10 monomers at each pole (apex). The overall assembly thus consists of 28 proteins, with units of cochaperonin Hsp10 sealing clients into the folding chamber (the ring cavity) where folding to the native state subsequently occurs.

Interactions between rings and Hsp10 caps are fundamental for the chaperone to achieve proper function. The simultaneous presence of distinct conformational states, also observed experimentally, may be considered necessary to engage Hsp10 and to adapt to the structural requirements of the maturing client.

#### Principal component analysis and conformational descriptors

Consistent with the larger degree of packing and conformational restraint determined by the presence of Hsp10 capping the double ring, PC1 and PC2 projections depict a much more rigid situation in **F** (Figure 8a). Constituent monomers of Hsp60 are locked in an even more open conformation and explore a much more limited portion of space compared to the previous cases (*cf.* triangular dots). It can be clearly seen that in all cases, either one very compact conformational state or at most two very close ones (PC1 at 100 and 110; PC2 always at 175) are retained throughout the entire simulation.

As in all other complexes examined so far, clear differences can be observed compared to **M**, with **δ** *vs.* **γ** graphs in Figure 8b showing a drastic change in both values for both variants and across protonation states: presence of substrate and cations in each active site brings aspartates (**δ**) clearly closer in **F**, apical opening angles **γ** transition from about 80° in **M** to either 120° or just below 120°—more open, in fact, than in **S** or **D**, and again close to the starting value (triangular dot; Figure 8b). Similarly to what happens for **D**, experiments suggest that once the **F** complexes are formed, **F_WT_** tends to be more active than **F_V72I_**, with a more pronounced production of active clients.^[14]^ While different distributions of **γ** values in our simulations for **F_WT_** and **F_V72I_** could provide a possible molecular explanation for this observation, we should take this with some caution as, consistently with greater conformational compactness, differences are rather less perceptible than those seen with **D_WT_** and **D_V72I_**. More specifically, in **F_WT_**, the apical domain opening tends to slightly exceed 120° degrees (PC1 peak at 110), while it more favourably samples values just under 120° (PC1 peak at 100) in **F_V72I_**, closer to the CryoEM structure. We again suggest a model whereby easier apical opening can be associated with greater volumes in the folding chamber to accommodate the client, and greater clearance when driving folding.

Even more intriguing trends emerge when focussing on Cγ distances in the catalytic aspartates (**δ**; Figure 8b), particularly when examined in conjunction with associated RDFs (Figures S7-S9). Indeed, if all these graphs are simultaneously compared with their counterparts for **D** (Figure 8; Figures S4-S6), one can see that the differences emerging between **D_WT_** and **D_V72I_** are replicated quite well for **F_WT_** *vs.* **F_V72I_**, even retaining those qualitative peculiarities that are distinctive of each protonation state, and clearly indicating that V72I has an impact on reactivity. (1) For Asp50-Asp397 (Figure S7), we again observe distances of around 5-5.5 Å, with a shoulder at around 4.5 Å that only exists in **F_V72I-AspAsp_** and not in **F_WT-AspAsp_**; with respect to **D**, we only note that the separation between the two peaks becomes more prominent, and the extra easiness with which the V72I variant is able to overcome the –2 charge is thus more evident. (2) For Asp50-Ash397 (Figure S9), we again observe the disappearance of all other peaks, and formation of a sharp peak at just over 4 Å that is statistically heavier in **F_V72I-AspAsh_**. (3) The Ash50-Asp397 case (Figure S8) is again the only one in which aspartates are able to approach each other more closely in the WT variant, and retain two peaks at 4.25 Å and 6 Å.

Once again, trends in **γ** are independent from trends in **δ**.

#### DF Analysis

Here, we again only focus on the P_avg_ projection of the ΔDF matrix for simulations of **F_WT_** and **F_V72I_**, which we show in Figure 5c for **F_WT-AspAsh_** *vs.* **F_V72I-AspAsh_**, and Figure S10 for the doubly deprotonated **F_WT-AspAsp_** *vs.* **F_V72I-AspAsp_**. In comparing P_avg_, there emerge some interesting observations that corroborate what we have seen considering P_avg_ for **M** and the lower complexes. While there is a clearly greater dependence on protonation states than in the other complexes, with a generalised loss of allosteric communication in doubly deprotonated Asp50-Asp397 (blue; Figure S10) that is not as prominent as in Asp50-Ash397 (closer to grey; Figure 5c), certain apical domains again consistently stand out, in orange, for a *gain* in allosteric prominence, particularly concentrated on helices α_9_ and α_10_, exactly as seen in other cases. A somewhat alternating orange-blue pattern is recognisable to some degree in both figures.

Further on the subject of protonation states, it is curious to see how the mutation impacts Hsp10 units in entirely opposite ways depending on whether Asp397 is protonated or not. In the former case (Figure 5c), and in marked contrast with the “alternating” allosteric profiles of apical domains, the mutation leads to increased allosteric communication with all Hsp10 units. In the latter case (Figure S10), Hsp10 units remain blue, signalling that the mutation is more effective in disrupting allosteric communication with Hsp10 units when both aspartates are deprotonated.

Importantly, these data reconfirm the fact that effects of the mutation extend pervasively over multiple scales, beyond individual monomers, right across even the largest possible complexes, with an impact on the organisation of the structural and dynamic states that underlie biological function. Overall, therefore, these results support a mechanistic model in which the mutation plays an active role in modifying the structural and conformational landscape of Hsp60, switching it to a more rigid and stable state at the expense of folding efficiency. Specifically, the mutation modifies the organisation of the active site, changing and perturbing the dynamics of the inter-domain dynamics, relaying the impact of the V72I mutation in each monomer to the folding chamber in which client remodelling takes place.

### V72I Rewires Allostery, Increasing Rigidity and Impacting Reactivity

Taken altogether, our cumulative data originating from MD simulations of **M**, **S**, **D**, and **F** are encouragingly consistent in describing a scenario in which mutation of Val72 to Ile significantly reshapes the allostery of individual Hsp60 monomers, with repercussions that affect all reaches of higher complexes. Data from simulations of **M** show that the mutation disrupts allosteric communication pathways present in WT Hsp60, which connect the equatorial and apical domains through the catalytically important intermediate domain: it does so by interrupting signals travelling across the adjacent equatorial helix α_18_ and the catalytically relevant α_14_. Instead, allosteric signals in mutant Hsp60 are ostensibly rewired so that the apical domain establishes more direct allosteric communication with certain interfacial regions of the equatorial domain: as a result, not only does it undergo some degree of internal rigidification, but it also loses its allosteric sensitivity to ATP hydrolysis, as is indeed explained below.

The greatest differences are of course seen ‘vertically’ *i.e.*, with increasing complex size, and are generally similar in both WT and mutant Hsp60, particularly when viewed in terms of apical opening (**γ**) and rotation (**χ**). The conformational freedom of **M** in WT and V72I Hsp60 is unsurprisingly unmatched in any of the other complexes, with individual monomers in **S**, **D**, and especially **F** clearly much more restricted in their movements, and apical domains constrained in increasingly open positions.

Yet, our data also provide interesting indications that the differences introduced by V72I itself systematically reverberate beyond isolated monomers. To summarise these mutation-related differences as concisely as possible, one could say that with the exception of **S**—where its effects are imperceptible—V72I is almost invariably seen to restrict the movement of apical domains so that they are prevented from looping through their conformational cycle (*i.e.*, dynamic states that are relevant for Hsp10 and client recognition become less easily accessible). We thus provide further proof that restriction of movement (rigidification) is what is behind the greater ease with which mutant complexes of Hsp60 can be isolated.^[14, 19]^

Apical domains in **M_V72I_** are seen to be able to rotate to the conformation they have in **S**, for example, but, unlike apical domains in **M_WT_**, they are visibly prevented from opening to reach the more raised conformation they should attain to get a functional **D_V72I_**. Out of all simulated complexes, monomers in **D_V72I_** are those that are most prominently and systematically affected by the mutation, and this time they are more favoured to stay open compared to unmutated monomers in **D_WT_**: this is true regardless of the protonation state of the catalytic dyad and, while not always synonymous with rigidification, there does remain a preference for open states compared to the wild-type double ring. Apical domains in **F_WT_** and **F_V72I_** are again much more restricted in their movement, but even in this constrained situation, there is a slight tendency for mutant apical domains to close back to the state they had in **D_V72I_**, prior to Hsp10 recruitment.

Throughout the higher complexes, the pattern of allosteric rewiring observed in **M** is generally replicated, but in a curiously nonuniform fashion. This is in line with the differential ‘up-down’ behaviour of apical domains previously highlighted in the literature.^[14]^ In any case, our simulations keep consistently suggesting that the disruption of allostery across the intermediate domain (and therefore across the catalytic dyad) is definitely affected by the mutation, suggesting that reactivity becomes decoupled from the fate of apical domains.

Having mentioned the catalytic dyad in Figure 1b as a key element that is impacted by V72I as soon as it becomes decoupled from apical domain opening / intermediate domain closing, we finally discuss the effects that the V72I mutation might have on reactivity according to our simulations, as suggested by Asp50:Cγ-Asp397:Cγ distances. As a reminder, Asp397 lies on intermediate domain helix α_14_, which is one of the elements undergoing the greatest loss of allosteric cross-communication upon mutation of Val72 to Ile. The issue of reactivity was copiously discussed in recent literature,^[14, 19]^ and experimentally addressed even in this work (*vide supra*), with increasing evidence that ATP hydrolysis rates increase in V72I-mutant Hsp60, especially in **D**. To briefly recap clues emerging from our own simulations of ATP-bound complexes **D** and **F**, we should focus on Asp50:Cγ-Asp397:Cγ plots (Figure 6b and 8b) and the associated RDFs (Figures S3-S5, S7-S9). All these plots unequivocally suggest that, regardless of protonation state, the aspartate dyad in **D_WT_** and **F_WT_** exhibits clearly different conformational dynamics / patterns than it does in **D_V72I_** and **F_V72I_**, and that the distribution in mutant Hsp60 complexes is indeed suggestive of augmented reactivity in the mutant (*vide infra*).

In particular, while in all systems with both deprotonated aspartates Cγ-Cγ distances never quite manage to get as short as 4 Å, it is intriguing to see that, in both **D_V72I-AspAsp_** and **F_V72I-AspAsp_**, there is a small but distinctive peak (‘shoulder’) at around 4.5 Å that is absent in RDFs of **D_WT-AspAsp_** and **F_WT-AspAsp_** (whose leftmost peak remains at just over 5 Å). This clearly indicates that mutant Hsp60 monomers in these complexes are somehow better equipped to overcome the significant energetic cost of bringing two negatively charged aspartates together (something that already occurs in the mutant CryoEM). As confirmed by high pK_a_ predictions (*cf. Computational Methods* section), bringing dyad aspartates close together renders them more basic, and likely better equipped to initiate ATP hydrolysis.

In systems where one of the two aspartates becomes protonated, on the other hand, the decrease in charge and formation of hydrogen bonds makes it possible for sidechains to approach each other more closely: this is clearly reflected in peaks generally shifting to the left. Even in this case, however, there is a matching distribution in mutant systems **D_V72I_** and **F_V72I_** that is distinctive from that in WT systems **D_WT_** and **F_WT_**. Curiously, simulations with protonated Asp50 indicate (higher RDF peaks) that it is easier for systems with WT chaperonin to keep the catalytic dyad closer together (*i.e.*, **D_WT-AshAsp_** and **F_WT-AshAsp_**); the opposite is true for systems in which Asp397 is protonated.

At any rate, since preliminary QM calculations (data not shown) indicate that the cost of exchanging a proton between Asp50 and Asp397 is merely in the order of 1 kcal mol^−1^, it is probably likelier that the actual conformational distribution of the aspartate dyad sidechains is somewhat smeared between these two ‘extremes’. Taken collectively, and postulating that closer, deprotonated aspartates might be more reactive, we can conclude that in complexes where the substrate is present in active sites alongside K^+^ and Mg^2+^ (*i.e.*, **D** and **F**), our simulations are indeed consistently suggestive that the V72I mutation might lead to increased ATPase activity. Further proving this point, however, naturally requires additional calculations.

Similarly, while “horizontal” increases in reactivity from **D_WT_** to **D_V72I_** and from **F_WT_** to **F_V72I_** are more readily identifiable by comparing RDFs, “vertical” cross-comparison (**D_WT_**/**F_WT_** and **D_V72I_**/**F_V72I_**) makes it less easy to identify the increase in wild-type reactivity upon addition of Hsp10, and the decrease in mutant reactivity. Again, further simulations are required to clarify this point.

### ATP Cleavage has an Immediate Effect on Crucial Allosteric Hotspots

To complete this part of the study, we also simulated **D** *post-*hydrolysis for both WT and mutant Hsp60 (Table 1; **D_WT-ADP-Pi_** and **D_V72I-ADP-Pi_**), by replacing ATP in each active site with ADP and [HPO_4_]^2–^ (P_i_), and protonating Asp397 (*i.e.*, treating it as the base driving hydrolysis). Comparison of **δ** *vs*. **γ** for **D_AspAsp_** and **D_ADP-Pi_**, in Figure 9a, unequivocally shows how the tendency after hydrolysis is to move towards an opening of the pocket that would favour nucleotide release from the active site after client folding, as shown by Cγ-Cγ distance **δ** clearly reapproaching **M** and **S** values (round dots; *i.e.*, aspartates returning farther away). Furthermore, the distribution of **γ** points to a relowering of the apical domains.

**Figure 9.**
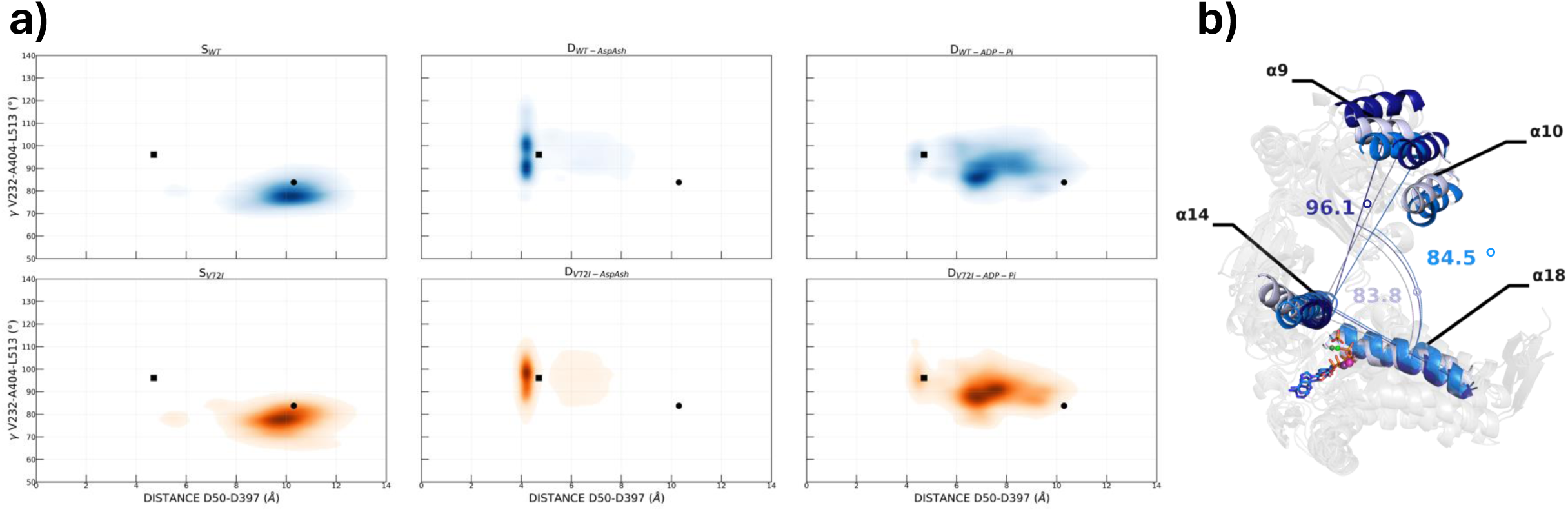
Motions in D_ADP-Pi_ during MD simulations. a) Distribution of distance **δ** and angle **γ** (see main text) in MD metatrajectories of **S** (leftmost panels), **D_AspAsh_** (middle panels), and **D_ADP-Pi_** (rightmost panels), featuring either WT Hsp60 (top panels; blue) or mutant Hsp60 (bottom panels; orange). Round and square dots represent CryoEM structure values as described in Figures 3, 4, 6, 8. As in Figures 3, 4, 6, 8, colour intensity is proportional to the probability density function, which is directly proportional to the number of MD frames with a given set of values. b) Salient 𝛼-helices (and other structural components) of Hsp60 monomers involved in allosteric changes. **D_WT_** features are represented in the darkest blue, **D_WT-ADP-Pi_** in middle blue (marine), and **S_WT_** in the lightest blue. For reference, we retain stick representations of ATP and ADP-P_i_ (C atoms in a scale of blues following the secondary structure pattern). Cations Mg^2+^ and K^+^ are represented as green and magenta spheres, respectively (following the same pattern, darker for **D_WT_** and lighter colour for **D_WT-ADP-Pi_**). Angles and labels follow the same pattern of color, being 83.8° **S_WT_**, 84.5° **D_WT-ADP-Pi_**, and 96.1° **D_WT_**.

To better illustrate our structural reasoning, we extrapolated representative (highly populated) conformations from **D_WT-ADP-Pi_** simulations to compare with the crystal structures of **S_WT_** and **D_WT_**. The four α-helices most prominently involved in allostery are highlighted in a scale of blues (Figure 9b). In line with biological expectations, α_14_ in **D_WT-ADP-Pi_** is revealed to occupy a position that is halfway between **D_WT_** and **S_WT_**, opening up with respect to the ‘reactive’ situation in **D_WT_** wherein α_14_ and α_18_ are more closely associated, and poised to facilitate substrate release. Similarly, it is clear from the reclosure of apical helices α_9_ and α_10_ how the first step in hydrolysis is sufficient to destabilise Hsp60-client interfaces in **D** and Hsp60-Hsp10 interfaces in **F** (and thus may well be conducive to client remodelling and **F** disassembly).

## Conclusions

Hsp60 is a multimeric chaperonin that, in combination with its cochaperone Hsp10, is instrumental in folding client proteins in the mitochondria. While the point mutation V72I is linked to autosomal dominant, SPG13 disease in humans and has dramatic effects on Hsp60 structure and function in vitro (Figure S1), its effects at the molecular level remain unclear. We considered this mutation to be an ideal biological model for asking this question because it preserves some aspects of chaperone function, such as modest amounts of client refolding, thus providing insight into a functional, but crippled, molecular machine.

In this work, with considerable computational effort, we have modelled and comparatively analysed 18 different complexes of Hsp60 (9 WT, 9 V72I mutants) with multimicrosecond long atomistic molecular dynamics simulations. Simulated systems range from the monomeric state **M** of Hsp60 all the way up to football-shaped complex **F**, which is an assembly of 28 different proteins (Hsp60_14_-Hsp10_14_).

To the best of our knowledge, this is the first comprehensive *in silico* investigation into the links between structure, dynamics and function of Hsp60, together with the elucidation of the allosteric impact of the V72I mutation, at such a different range of scales. A particularly important point emerging from our work is the direct link we can make with the most recent findings on Hsp60 complexes,^[14, 19]^ including experimental data presented here along with the simulations.

Combining a series of different allostery detection methods and analyses, our simulations unveil the mechanisms through which one single mutation at a protein site distal from the reactive centre of Hsp60 is able to influence conformational dynamics of the protein at multiple scales, affecting the selection of specific dynamic states from the monomer state to large scale multiprotein assemblies. In the monomer, mutation V72I modifies the internal organisation of functional dynamics and the mechanisms of coordination among distal sets of residues, disrupting allosteric signals that transit across the intermediate domain while rigidifying the substructures (such as β-strands) that form the inter-monomer interfaces in the larger assemblies, and establishing a more direct equatorial-apical dialogue. This preorganisation can better poise the system for the formation of larger aggregates and supports a molecular model for the experimentally observed hyperstability of higher Hsp60 V72I oligomers.

Moreover, we find that effects observed in the monomer extend to the larger supramolecular complexes. Interestingly, particularly in the double ring complexes **D**, we consistently observe a general rigidification upon mutation: this is generally consistent with the observed stabilisation that ultimately favoured their structural resolution via Cryo-EM.^[14]^

Importantly, the mutant shows a clear impact on the allosteric fingerprint of the apical domains in the assembly. In this context, while we cannot observe full-scale, reversible and alternate motions of apical domains, our simulations fully delineate the nonuniform microscopic changes in dynamics introduced by V72I that are plausibly conducive to alterations in macroscopic apical domain movement. These alterations are those that so clearly usher in hyperstabilisation, impaired biological function, and diminished cochaperonin/client recognition, due to a shift in conformational ensembles populated by the domains. Furthermore, observed perturbations in the dynamics can in all likelihood be linked to the ‘up’-‘down’ movement inferable from structures published by Braxton *et al.*^[14]^ This is achieved through the local disruption and reassembly of interactions that will eventually result in structural deformations (see ^[46]^) on longer time scales.

In this context, our results also highlight how the higher rigidity of α_9_ and α_10_ dedicated to interactions with clients and cochaperones, combined with a lower general capacity of conveying allosteric signals coming from the equatorial and intermediate ATPase domains, cause the partial impairment in the chaperone functions, in line with experimental observations. While decoupled allosterically from the apical domains, a preliminary interpretation of our simulations based on important differences in the catalytic Asp50-Asp397 dyad upon mutation suggests a slight increase in ATPase activity: this is again in line with previous experimental observations^[14, 19–20, 45]^ and with the experimental data presented here.

From these results, we propose a model where direct coupling of mutation-induced dynamic perturbation in the monomeric protein with active site organisation and conformational selection of domain organisation reverberates at multiple scales in the rearrangement of the functionally oriented dynamics of diverse large multiprotein assemblies, providing an atomistic picture of how even a single mutation can create complexes with altered functions compared to the native sequence.

## Supporting information

Textual Supporting Information

Input scripts and videos

## Acknowledgements

This work was supported by grants from Associazione Italiana Ricerca sul Cancro (AIRC) under IG 2022 - ID. 27139; PNRR-MUR Research Programme CN00000013 “National Centre for HPC, Big Data and Quantum Computing”; PRIN grant 20209KYCH9; from IMMUNO-HUB, T4-CN-02, from the Ministero della Salute (Italy). F.G. is supported by the AIRC Individual Fellowship Love Design 2021 (ID 26647-2021). Authors wish to thank Prof. Sílvia Osuna (University of Girona) for her precious assistance in obtaining the *DynaComm.py* code.

## Electronic Supporting Information

MD preproduction details — Derivation of SPM matrices — Biochemical methods — Native PAGE results — Additional DF analyses — Asp50:Cγ–Asp397:Cγ RDF plots (PDF)

MD: input scripts, coordinates, topologies — SPM: input scripts, matrices and output — PCA: input scripts — Movies of PC1 and PC2 — Movies of the ΔDF matrix (projections P_0_ to P_525_) (Drive folder: https://drive.google.com/drive/folders/1IqlLigmQdIOWswc-rUciF7cL1oOqE8Gs?usp=sharing)

## References

[1] a) M. Y. Cheng, F. U. Hartl, J. Martin, R. A. Pollock, F. Kalousek, W. Neuper, E. M. Hallberg, R. L. Hallberg, A. L. Horwich, Nature 1989, 337 620–625; b) Y. E. Kim, M. S. Hipp, A. Bracher, M. Hayer-Hartl, F. U. Hartl, Annu. Rev. Biochem. 2013, 2013, 323–355.

[2] a) J. C. Ghosh, T. Dohi, H. K. Byoung, D. C. Altieri, J. Biol. Chem. 2008, 283 5188–5194; b) X. S. Li, Q. Xu, X. Y. Fu, W. S. Luo, PLoS One 2014, *9* e107507.

[3] K. C. Walls, P. Coskun, J. L. Gallegos-Perez, N. Zadourian, K. Freude, S. Rasool, M. Blurton-Jones, K. N. Green, F. M. LaFerla, Journal of Biological Chemistry 2012, 287, 30317–30327.

[4] F. U. Hartl, Nature 1996, 381, 571–580.

[5] a) D. Balchin, M. Hayer-Hartl, F. U. Hartl, Science 2008, 353, 6294; b) B. Bukau, A. L. Horwich, Cell 1998, 92 351–366.

[6] Z. Xu, A. L. Horwich, P. B. Sigler, Nature 1997, 388, 741–750.

[7] H. S. Rye, S. G. Burston, W. A. Fenton, J. M. Beechem, Z. Xu, P. B. Sigler, A. L. Horwich, Nature 1997, 388, 792–798.

[8] a) A. L. Horwich, G. W. Farr, W. A. Fenton, Chem. Rev. 2006, 106, 1917–1930; b) J. Ma, P. B. Sigler, Z. Xu, M. Karplus, J. Mol. Biol. 2000, 302, 303–313; c) J. Weaver, M. Jiang, A. Roth, J. Puchalla, J. Zhang, H. S. Rye, Nat. Commun. 2017, 8, 15934; d) C. L. Hyeon, G. H.; Thirumalai, D., Proc. Natl. Acad. Sci. U. S. A. 2006, 103, 18939–18944; e) Y.-C. Tang, H.-C. Chang, A. Roeben, D. Wischnewski, N. Wischnewski, M. J. Kerner, F. U. Hartl, M. Hayer-Hartl, Cell 2006, 125, 903–914.

[9] a) S. Nisemblat, O. Yaniv, A. Parnas, F. Frolow, A. Azem, Proc. Natl. Acad. Sci. U. S. A. 2015, 112, 6044–6049; b) Y. Gomez-Llorente, F. Jebara, M. Patra, R. Malik, S. Nisemblat, O. Chomsky-Hecht, A. Parnas, A. Azem, J. A. Hirsch, I. Ubarretxena-Belandia, Nature Communications 2020, 11, 1916; c) D. P. Klebl, M. C. Feasey, E. L. Hesketh, N. A. Ranson, H. Wurdak, F. Sobott, R. S. Bon, S. P. Muench, iScience 2021, 24, 102022; d) J. C.-Y. Wang, L. Chen, Scientific Reports 2021, 11, 14809.

[10] a) G. B. Levy-Rimler, R. E.; Ben-Tal, N.; Azem, A., FEBS Lett. 2002, 529, 1–5; b) C. Weiss, F. Jebara, S. Nisemblat, A. Azem, Frontiers in Molecular Biosciences 2016, 3.

[11] P. V. Viitanen, G. Lorimer, W. Bergmeier, C. Weiss, M. Kessel, P. Goloubinoe, Methods Enzymol. 1998, 290, 203–217.

[12] H. R. Saibil, W. A. Fenton, D. K. Clare, A. L. Horwich, J. Mol. Biol. 2013, 425, 1476– 1487.

[13] G. Levy-Rimler, P. Viitanen, C. Weiss, R. Sharkia, A. Greenberg, A. Niv, A. Lustig, Y. Delarea, A. Azem, European Journal of Biochemistry 2001, 268, 3465–3472.

[14] J. R. Braxton, H. Shao, E. Tse, J. E. Gestwicki, D. R. Southworth, Nature Structural & Molecular Biology 2024.

[15] M. J. Clie, C. Limpkin, A. Cameron, S. G. Burston, A. R. Clarke, Journal of Biological Chemistry 2006, 281, 21266–21275.

[16] K. L. Nielsen, N. J. Cowan, Molecular Cell 1998, 2, 93–99.

[17] E. S. Bochkareva, N. M. Lissin, G. C. Flynn, J. E. Rothman, A. S. Girshovich, Journal of Biological Chemistry 1992, 267, 6796–6800.

[18] a) A. Parnas, M. Nadler, S. Nisemblat, A. Horovitz, H. Mandel, A. Azem, Journal of Biological Chemistry 2009, 284, 28198–28203; b) C. Cömert, L. Brick, D. Ang, J. Palmfeldt, B. F. Meaney, M. Kozenko, C. Georgopoulos, P. Fernandez-Guerra, P. Bross, Cold Spring Harb. Mol. Case Stud. 2020, 6, 1–13; c) D. Magen, C. Georgopoulos, P. Bross, D. Ang, Y. Segev, D. Goldsher, A. Nemirovski, E. Shahar, S. Ravid, A. Luder, B. Heno, R. Gershoni-Baruch, K. Skorecki, H. Mandel, The American Journal of Human Genetics 2008, 83, 30–42; d) J. J. Hansen, A. Dürr, I. Cournu-Rebeix, C. Georgopoulos, D. Ang, M. N. Nielsen, C.-S. Davoine, A. Brice, B. Fontaine, N. Gregersen, P. Bross, The American Journal of Human Genetics 2002, 70, 1328-1332.

[19] A. Syed, J. Zhai, B. Guo, Y. Zhao, J. C.-Y. Wang, L. Chen, Structure 2024, 32, 575–584.e573.

[20] P. Bross, S. Naundrup, J. Hansen, M. N. Nielsen, J. H. Christensen, M. Kruhøeer, J. Palmfeldt, T. J. Corydon, N. Gregersen, D. Ang, C. Georgopoulos, K. L. Nielsen, Journal of Biological Chemistry 2008, 283, 15694–15700.

[21] Schrödinger LTD, 2015.

[22] D. A. Case, H. M. Aktulga, K. Belfon, D. S. Cerutti, G. A. Cisneros, V. W. D. Cruzeiro, N. Forouzesh, T. J. Giese, A. W. Götz, H. Gohlke, S. Izadi, K. Kasavajhala, M. C. Kaymak, E. King, T. Kurtzman, T.-S. Lee, P. Li, J. Liu, T. Luchko, R. Luo, M. Manathunga, M. R. Machado, H. M. Nguyen, K. A. O’Hearn, A. V. Onufriev, F. Pan, S. Pantano, R. Qi, A. Rahnamoun, A. Risheh, S. Schott-Verdugo, A. Shajan, J. Swails, J. Wang, H. Wei, X. Wu, Y. Wu, S. Zhang, S. Zhao, Q. Zhu, T. E. Cheatham, III, D. R. Roe, A. Roitberg, C. Simmerling, D. M. York, M. C. Nagan, K. M. Merz, Jr., Journal of Chemical Information and Modeling 2023, 63, 6183–6191.

[23] M. H. M. Olsson, C. R. Søndergaard, M. Rostkowski, J. H. Jensen, J. Chem. Theory Comput. 2011, 7, 525–537.

[24] A. Koike-Takeshita, K. Mitsuoka, H. Taguchi, Journal of Biological Chemistry 2014, 289, 30005–30011.

[25] J. Wang, D. C. Boisvert, J. Mol. Biol. 2003, 3, 843–855.

[26] J. A. Maier, C. Martinez, K. Kasavajhala, L. Wickstrom, K. E. Hauser, C. Simmerling, Journal of Chemical Theory and Computation 2015, 11, 3696–3713.

[27] W. L. Jorgensen, J. Chandrasekhar, J. Madura, R. W. Impey, M. L. Klein, J. Chem. Phys. 1983, 79, 926–935.

[28] I. S. Joung, T. E. Cheatham, J. Phys. Chem. B 2008, 112, 9020–9041.

[29] O. Allnér, L. Nilsson, A. Villa, J. Chem. Theory Comput. 2012, 8, 1493–1502.

[30] K. L. Meagher, L. T. Redman, H. A. Carlson, J. Comput. Chem. 2003, 24, 1016–1025.

[31] S. Kashefolgheta, A. Vila Verde, Phys. Chem. Chem. Phys. 2017, 19, 20593–20607.

[32] D. A. Case, T. E. Cheatham Iii, T. Darden, H. Gohlke, R. Luo, K. M. Merz Jr, A. Onufriev, C. Simmerling, B. Wang, R. J. Woods, Journal of Computational Chemistry 2005, 26, 1668–1688.

[33] R. Salomon-Ferrer, A. W. Götz, D. Poole, S. Le Grand, R. C. Walker, J. Chem. Theory Comput. 2013, 9, 3878–3888.

[34] R. J. Loncharich, B. R. Brooks, R. W. Pastor, Biopolymers 1992, 32, 523–535.

[35] H. J. C. Berendsen, J. P. M. Postma, W. F. van Gunsteren, A. Di Nola, J. R. Haak, J. Chem. Phys. 1984, 81, 3684–3690.

[36] T. Darden, D. York, L. Pedersen, J. Chem. Phys. 1993, 98.

[37] S. Miyamoto, P. A. Kollman, J. Comp. Chem. 1992, 13, 952–962.

[38] M. L. Waskom, J. Open Source Softw. 2021, 6, 3021.

[39] a) M. Castelli, A. Magni, G. Bonollo, S. Pavoni, F. Frigerio, A. S. F. Oliveira, F. Cinquini, S. A. Serapian, G. Colombo, Protein Science 2024, 33, e4880; b) M. Castelli, F. Marchetti, S. Osuna, A. S. F. Oliveira, A. J. Mulholland, S. A. Serapian, G. Colombo, J. Am. Chem. Soc. 2024, 146, 901–919; c) G. Morra, R. Potestio, C. Micheletti, G. Colombo, Plos Comput. Biol. 2012, 8, e1002433; d) E. Moroni, D. A. Agard, G. Colombo, J. Chem. Theory Comput. 2018, 14, 1033–1044.

[40] G. Colombo, Curr. Opin. Struct. Biol. 2023, 83, 102702.

[41] a) A. Romero-Rivera, M. Garcia-Borràs, S. Osuna, ACS Catal. 2017, 7, 8524–8532; b) S. Osuna, Wiley Interdiscip. Rev.: Comput. Mol. Sci. 2021, 11, e1502.

[42] P. Pons, M. Latapy, Journal Graph Algorithms Appl. 2006, 10, 191–218.

[43] G. Casadevall, J. Casadevall, C. Duran, S. Osuna, Protein Engineering, Design and Selection 2024, 37, gzae005.

[44] A. Amadei, Linssen, A. B. M., Berendsen, H. J. C., Proteins: Struct. Funct. Genet. 1993, 17, 412–425.

[45] L. Chen, A. Syed, A. Balaji, Scientific Reports 2022, 12, 18321.

[46] a) K. A. Henzler-Wildman, M. Lei, V. Thai, S. J. Kerns, M. Karplus, D. Kern, Nature 2007, 450, 913–918; b) K. A. Henzler-Wildman, V. Thai, M. Lei, M. Ott, M. Wolf-Watz, T. Fenn, E. Pozharski, M. A. Wilson, G. A. Petsko, M. Karplus, C. G. Hubner, D. Kern, Nature 2007, 450, 838–844; c) F. Pontiggia, D. Pachov, M. Clarkson, J. Villali, M. Hagan, V. Pande, D. Kern, Nat. Commun. 2015, 6, 7284; d) R. Nussinov, C.-J. Tsai, H. Jang, PLoS computational biology 2019, 15, e1006648–e1006648; e) S. J. Wodak, E. Paci, N. V. Dokholyan, I. N. Berezovsky, A. Horovitz, J. Li, V. J. Hilser, I. Bahar, J. Karanicolas, G. Stock, P. Hamm, R. H. Stote, J. Eberhardt, Y. Chebaro, A. Dejaegere, M. Cecchini, J.-P. Changeux, P. G. Bolhuis, J. Vreede, P. Faccioli, S. Orioli, R. Ravasio, L. Yan, C. Brito, M. Wyart, P. Gkeka, I. Rivalta, G. Palermo, J. A. McCammon, J. Panecka-Hofman, R. C. Wade, A. Di Pizio, M. Y. Niv, R. Nussinov, C.-J. Tsai, H. Jang, D. Padhorny, D. Kozakov, T. McLeish, Structure 2019, 27, 566–578; f) R. Nussinov, C.-J. Tsai, H. Jang, Current Opinion in Structural Biology 2021, 71, 43–50; g) R. Nussinov, M. Zhang, R. Maloney, Y. Liu, C.-J. Tsai, H. Jang, Journal of Molecular Biology 2022, 434, 167569; h) D. U. Ferreiro, E. A. Komives, P. G. Wolynes, Current Opinion in Structural Biology 2018, 48, 68–73; i) J. Chen, N. P. Schafer, P. G. Wolynes, C. Clementi, The Journal of Physical Chemistry B 2019, 123, 4497–4504.

