## Supplementary material for "How a pathogenic mutation impairs Hsp60 functional dynamics from monomeric to fully assembled states": Textual Supporting Information

#### MD preproduction details

Below is an overview of the conditions adopted for the preproduction stage in all of our MD replicas, which comprised two minimisation stages (300 + 300 steps), a solvent equilibration stage (9 ps), a restrained heating stage (20 ps), and an equilibration stage (2.040 ns), for a total of 2.069 ns. Unless otherwise stated, conditions were identical to the production stage (see main text).

##### Minimisation

Minimisation was carried out with the *sander* utility[1] and involved two stages, each comprising up to 300 minimisation steps: 10 using the steepest descent algorithm, and up to 290 using the conjugate gradient approach. The first stage was very conservative, with harmonic position restraints ($k$ = 5.0 kcal mol$^{-1}$ Å$^{-2}$) applied to all atoms except protein Hα. For the subsequent stage, however, restraints were entirely lifted. Neither stage entailed periodic boundary conditions.

##### Solvent Equilibration

Solvent Equilibration took place for 9 ps in the *NVT* ensemble, again using the *sander* utility,[1] henceforth introducing periodic boundary conditions. All atoms randomly assigned velocities matching a temperature of 25 K. Non-water atoms were positionally restrained ($k$ = 10.0 kcal mol$^{-1}$ Å$^{-2}$), whereas water atoms were allowed to move using a (shorter) timestep of 1 fs since, only at this stage, we did not impose any *SHAKE* constraints.[2] Also unlike subsequent stages, temperatures were enforced using Berendsen's thermostat.[3] Rapid solvent heating from 25 K to 400 K was imposed over the first 3 ns, with a rather stringent coupling constant of 0.2 ps. For the next 3 ns, the solvent was kept at 400 K, and for the final 3 ns, it was more gently cooled back to 25 K with a blander 2.0 ps coupling constant.

##### Restrained Heating to 300 K

Switching to *pmemd.cuda*[4] and to the Langevin thermostat[5] but with a more stringent 0.75 ps$^{-1}$ collision frequency to ensure quicker heating, restrained heating from 25 K to 300 K was carried out in the *NVT* ensemble for 20 ps. Specifically, harmonic position restraints are only applied to protein Cα atoms ($k$ = 5.0 kcal mol$^{-1}$ Å$^{-2}$): all other atoms are allowed to move. Atomic velocities were *not* retained from the previous stage, but randomly reassigned anew for each replica: subsequent stages, however, inherited atomic velocities issuing from the previous stage.

##### Equilibration

Equilibration takes place over 2.040 ns, switching to the *NpT* ensemble, wherein the 1.0 bar pressure is imposed exactly as during the production stage.[3] With respect to restrained heating, coupling to the Langevin thermostat is immediately softened[5] with a collision frequency of 1.0 ps$^{-1}$, and restraints on Cα atoms are gradually lifted. More specifically,restraints are lowered to $k$ = 3.75 kcal mol$^{-1}$ Å$^{-2}$ for the first 20 ps, further to $k$ = 1.75 kcal mol$^{-1}$ Å$^{-2}$ for the next 20 ps, and lifted entirely for the final 2 ns.

#### Derivation of Matrices for the SPM

As stated in the main text, derivation of the SPM for $\mathbf{M}_{\mathbf{WT}}$ and $\mathbf{M}_{\mathbf{V72I}}$ required the prior calculation, for each system, of an average pairwise distance matrix **d** and an average pairwise correlation matrix **C**. Derivation of **C** and **d** from 2 μs metatrajectories is straightforwardly achievable with the *cpptraj* utility:[6] we provide all input scripts electronically, as well as **C** and **r** for both systems. In particular, as *per* the formula for **C** elements $C_{ij}$ reported in the main text, calculation of **C** requires a reference structure: we take this to be the structure closest to the centroid of the most populated conformational cluster in each metatrajectory.

More specifically, reference structures (closest to cluster centroids) for $\mathbf{M}_{WT}$ and $\mathbf{M}_{V72I}$ are derived as follows:

- All metatrajectory frames are aligned to their very first frame (formerly the first frame in the production stage of replica 1), by minimising the root-mean-square deviation (RMSD) of all Cα atoms.
- An average structure is derived from this realigned metatrajectory, using the `average` command in *cpptraj*.[6]
- All metatrajectory frames are realigned to this average structure, again based on minimising the RMSD of all Cα atoms.
- Clustering is performed on these realigned metatrajectories after dropping (sieving) every other frame and thus reducing them to 20000 frames. Clustering is performed using the hierarchical agglomeration method as implemented in *cpptraj*,[6] and stopped once the number of clusters identified has dropped to 5. The representative structure to derive **C** is the one closest to the centroid of the most populated cluster.

Once reference structures are obtained, **C** matrices are derived as follows:

- Full metatrajectories (40000 frames each) are realigned for the third time, this time to their respective reference structure, again minimising the RMSD of all Cα atoms.
- The `matrix` command in *cpptraj*[6] is used to directly output **C**.

Similarly, **d** matrices are also derived with the `matrix` command in *cpptraj*[6] (exact syntaxes are extractable from the input scripts provided). **C** and **d** matrices derived for $\mathbf{M}_{WT}$ and $\mathbf{M}_{V72I}$ are passed as-is to the *DynaComm.py* code.[7]

### Biochemical Methods

#### Protein purification

Recombinant, human Hsp60 and its variants were expressed as previously described[8] (ref *DOI: 10.1039/d0ob00928h*). Briefly, *E. coli* BL21 (DE3) cells in terrific broth (TB; 1L) were grown at 37 °C with shaking to 0.6 $OD_{600}$, followed by induction with isopropyl β-D-1-thiogalactopyranoside (IPTG; final concentration 300 µM). After another 4 hours, cell pellets were generated and then re-suspended in His-binding buffer (50 mM TRIS, 10 mM Imidazole, 500 mM NaCl, pH 8) supplemented with protease inhibitors. Cells were lysed by sonication, pelleted by centrifugation, and the supernatant was incubated with Ni-NTA His-Bind Resin for 1 hour at 4 °C. The resin was washed with His-binding buffer, followed by His-washing buffer (50 mM TRIS, 30 mM imidazole, 300 mM NaCl, pH 8) and elution with His-elution buffer (50 mM TRIS, 300 mM Imidazole, 300 mM NaCl, pH 8). The N-terminal His-tags were removed by incubating the protein and TEV protease with 1 mM DTT for 4 hours at room temperature, followed by dialysis in 4 L of SEC buffer (50 mM TRIS, 0.3 M NaCl, 10 mM $MgCl_2$, pH 7.7) overnight.

#### Native gels

An aliquot of Hsp60 (5 µM; 15 µL) in sample buffer (5X: 0.05% Ponceau Red, 50% Glycerol, 250 mM 6- aminohexanoic acid, 50 mM Bis-Tris, pH 7.0) was loaded into precast 4–16% Novex NativePAGE Bis-Tris gels. Electrophoresis was performed at 150 V constant voltage for 4 hours at room temperature with 50 mM Bis Tris (pH 7.0) as anode buffer and 15 mM Bis Tris and 50 mM Tricine (pH 7.0) as cathode buffer. Gel was stained by Coomassie G-250 and scanned using ChemiDoc Touch Imager.

#### ATPase assays

ATPase activity was measured by malachite green assays, previously described[9]. To each well of a 96-well plate, Hsp60 solution (20 µL; 2 µM final) in assay buffer (100 mM Tris, 20 mM KCl, 10 mM $MgCl_2$, 0.017% Triton, pH 7.4) was added. Reactions were initiated by adding ATP (5 µL; 1 mM final) and incubated at 37 °C. Reactions were quenched by the addition of malachite green reagent (80 µL) which was added into each well, followed by 32% sodium citrate (10 µL). The plate was incubated at 37 °C for 15 min before measuring $OD_{620}$ on a SpectraMax M5 microplate reader (Molecular Devices, Sunnyvale, CA).

#### MDH refolding

Briefly, malate dehydrogenase (MDH, Sigma-10127256001) was denatured with guanidine buffer (7 M guanidine, 200 mM Tris, pH 7.4) for 1 h at room temperature. A sample of the denatured MDH (120 nM) was added to Hsp60 (3.33 µM) and Hsp10 (6.67 µM) in refolding buffer (100 mM Tris pH 7.4, 20 mM KCl, 10 mM $MgCl_2$, and 1 mM DTT). Then, 30 µL of this solution was dispensed into clear 384-well plates (CORNING, 3702) and ATP was added (20 µL of 2.5 mM). The final conditions were: 2 µM Hsp60, 4 µM Hsp10, 72 nM denatured MDH, 1 mM ATP). At each time point, the reaction was quenched by adding 10 µL of 500 mM EDTA and NADH absorbance at 340 nM was measured for 60 minutes in assay buffer (20 mM sodium mesoxalate monohydrate and 2.4 mM NADH in reaction buffer, Sigma-Aldrich). $A_{340}$ measurements were recorded at $T_0$ and then the time point at which 90% of substrate was consumed was reported. The $IC_{50}$ values were calculated using GraphPad Prism.

#### Native PAGE shows that $Hsp60^{V72I}$ is a stable oligomer

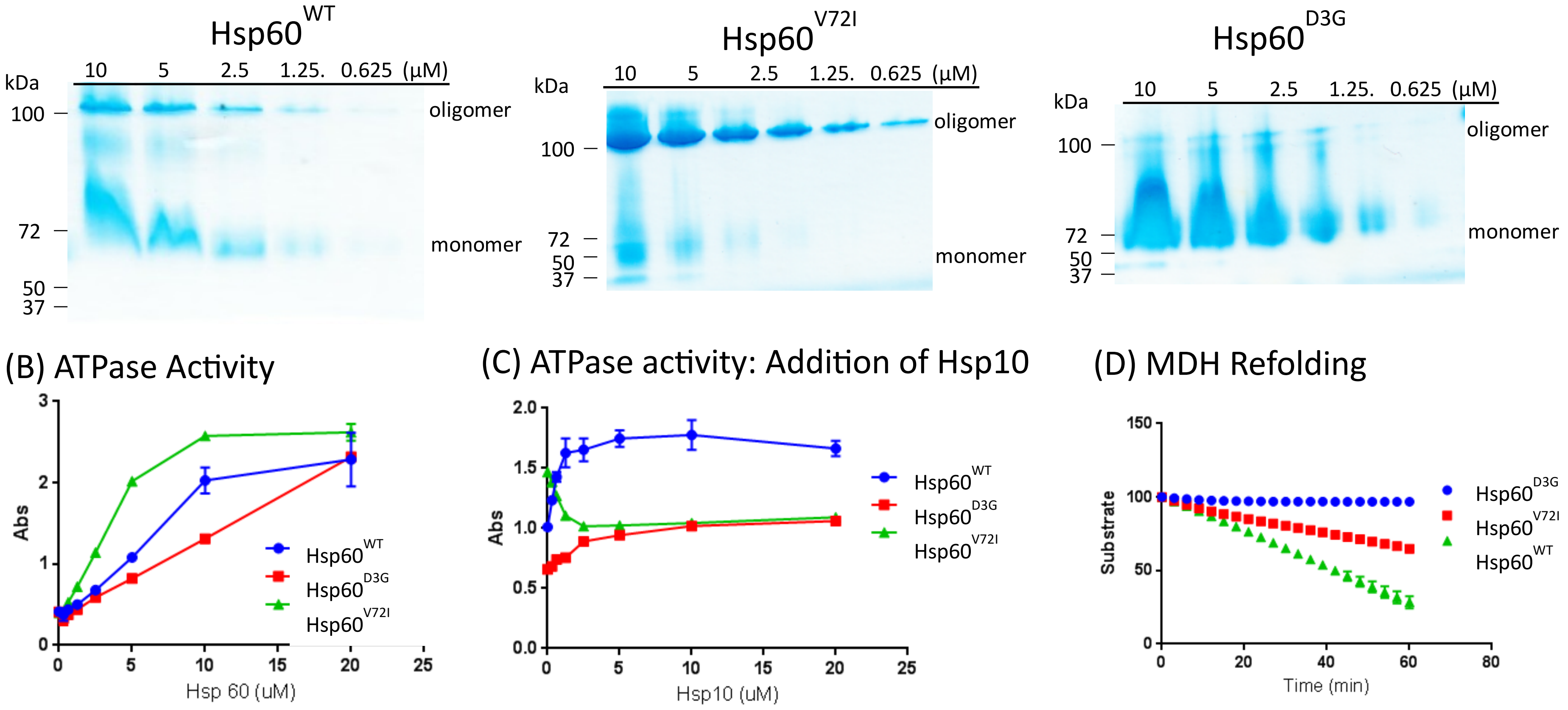

**Figure S1.** Biochemical characterisation of Hsp60WT and Hsp60V72I confirms a dramatic effect on ATPase activity and oligomer formation. (A) Native PAGE on purified, human Hsp60 and its variants. Wild type Hsp60 ($Hsp60^{WT}$) forms unstable oligomers that sample monomer, while $Hsp60^{V21I}$ forms stable oligomers at a range of concentrations. Results are representative of at least three replicates. (B) Steady state ATPase activity, as measured by malachite green assays. Results are the average of independent triplicates and the error bars represent standard deviation (SD). (C) ATPase assays of Hsp60 (2 µM) in the presence of Hsp10. Results are the average of independent triplicates and the error bars represent standard deviation (SD). (D) Refolding of malate dehydrogenase (MDH), as measured by disappearance of its colorimetric substrate (see Methods).

#### DF Analysis on MD Simulations of $M_{WT}$ and $M_{V72I}$

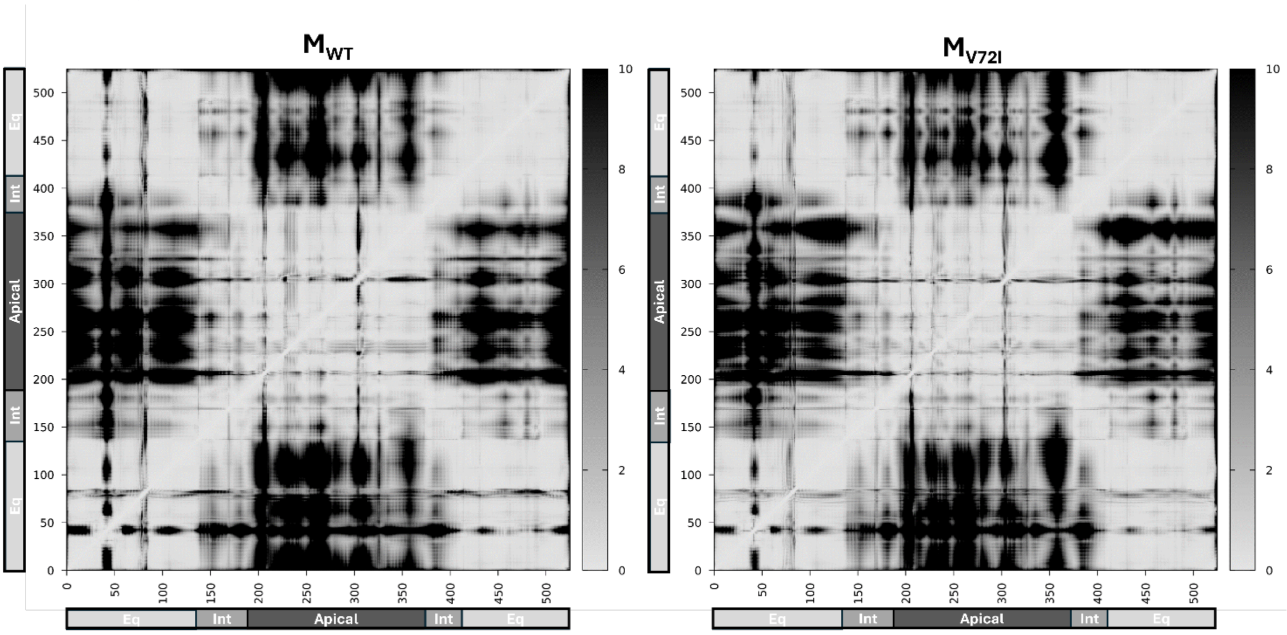

**Figure S2.** DF matrices calculated using Cα atoms of each residue pair in MD metatrajectories of $\mathbf{M_{WT}}$ and $\mathbf{M_{V72I}}$ (see Computational Methods section). DF scores are represented as depicted in the colour bar to the right of each matrix. Residue numbering is shown on both axes together with the corresponding domains: Equatorial (residues 1-133,410-525); Intermediate (residues 134-188,374-409); Apical (residues 189-373).

#### RDF Plots from MD Simulations of $D_{WT}$ and $D_{V72I}$

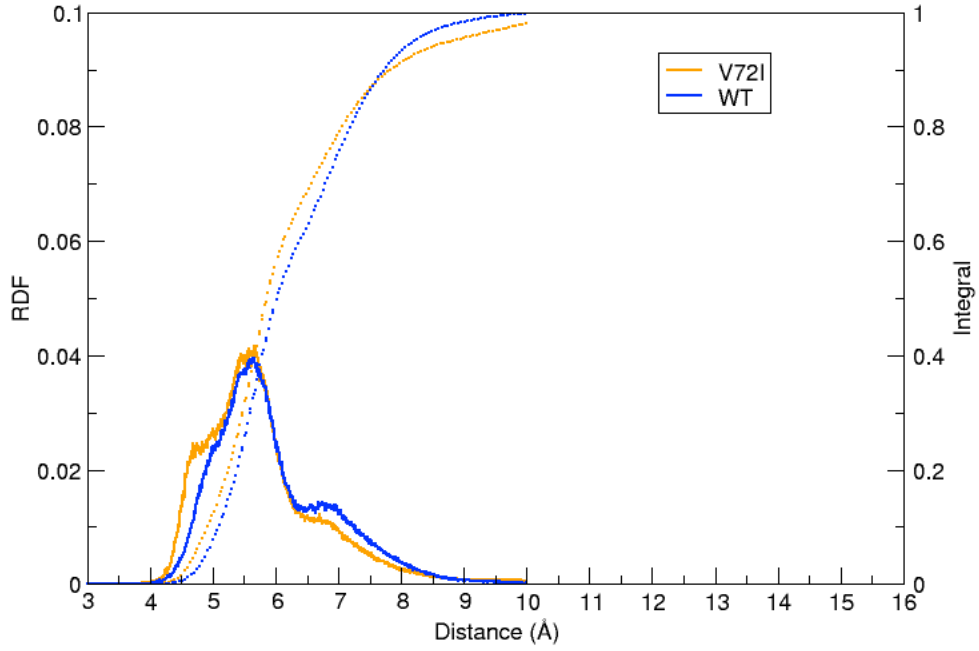

**Figure S3.** RDF plots of the Asp50:Cγ–Asp397:Cγ distance derived from MD metatrajectories of $\mathbf{D_{WT\text{-}AspAsp}}$ (blue) and $\mathbf{D_{V72I\text{-}AspAsp}}$ (orange). Dotted lines represent RDF integrals (right *y*-axis).

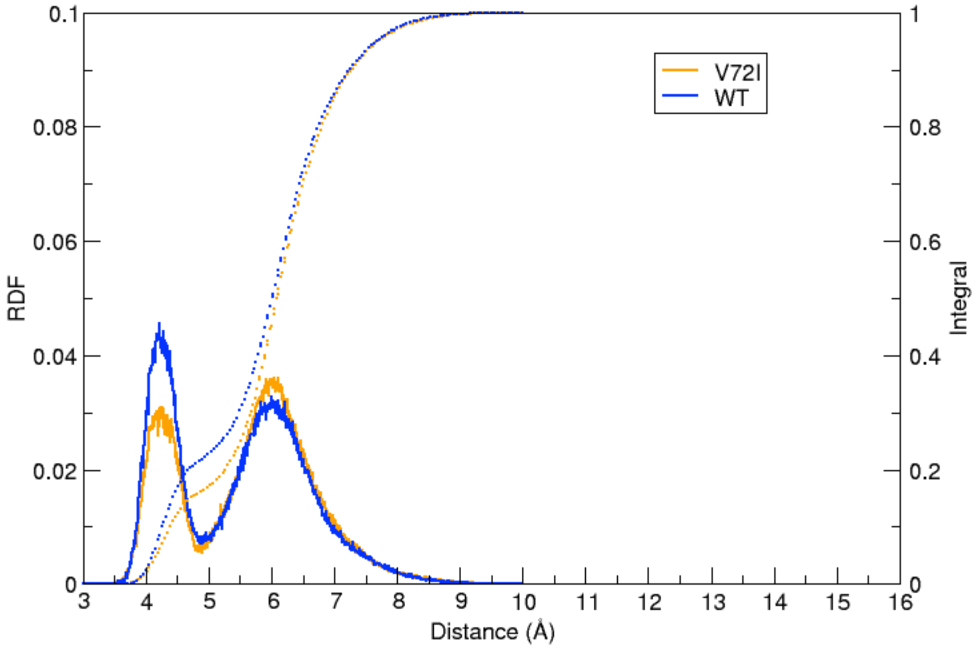

**Figure S4.** RDF plots of the Ash50:Cγ–Asp397:Cγ distance derived from MD metatrajectories of $\mathbf{D}_{\mathbf{WT\text{-}AshAsp}}$ (blue) and $\mathbf{D}_{\mathbf{V72I\text{-}AshAsp}}$ (orange). Dotted lines represent RDF integrals (right *y*-axis).

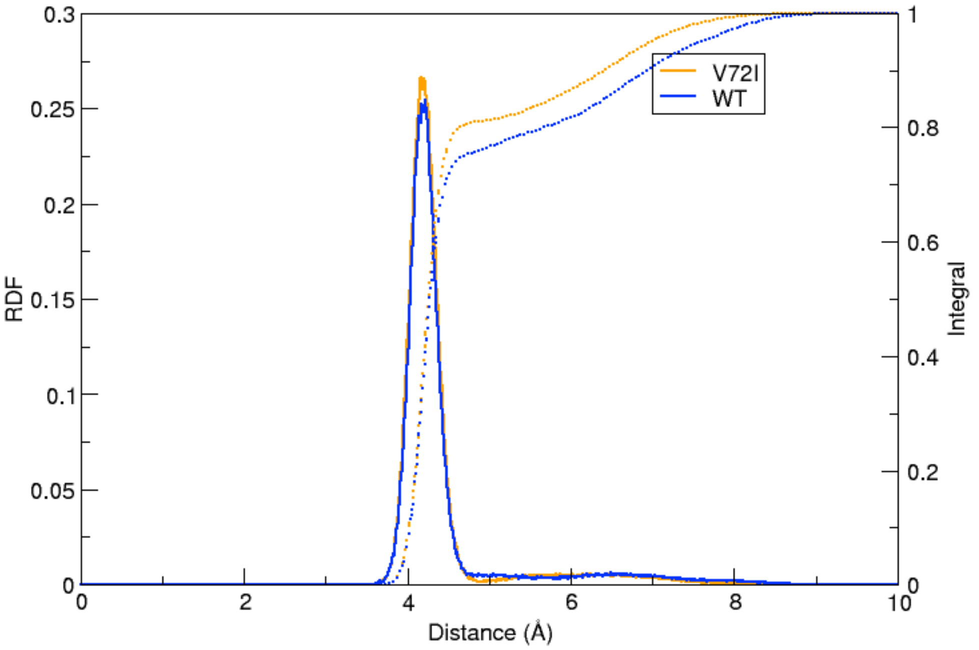

**Figure S5.** RDF plots of the Asp50:Cγ–Ash397:Cγ distance derived from MD metatrajectories of $\mathbf{D}_{\mathbf{WT\text{-}AspAsh}}$ (blue) and $\mathbf{D}_{\mathbf{V72I\text{-}AspAsh}}$ (orange). Dotted lines represent RDF integrals (right *y*-axis).

#### DF Analysis on MD Simulations of $D_{WT}$ and $D_{V72I}$

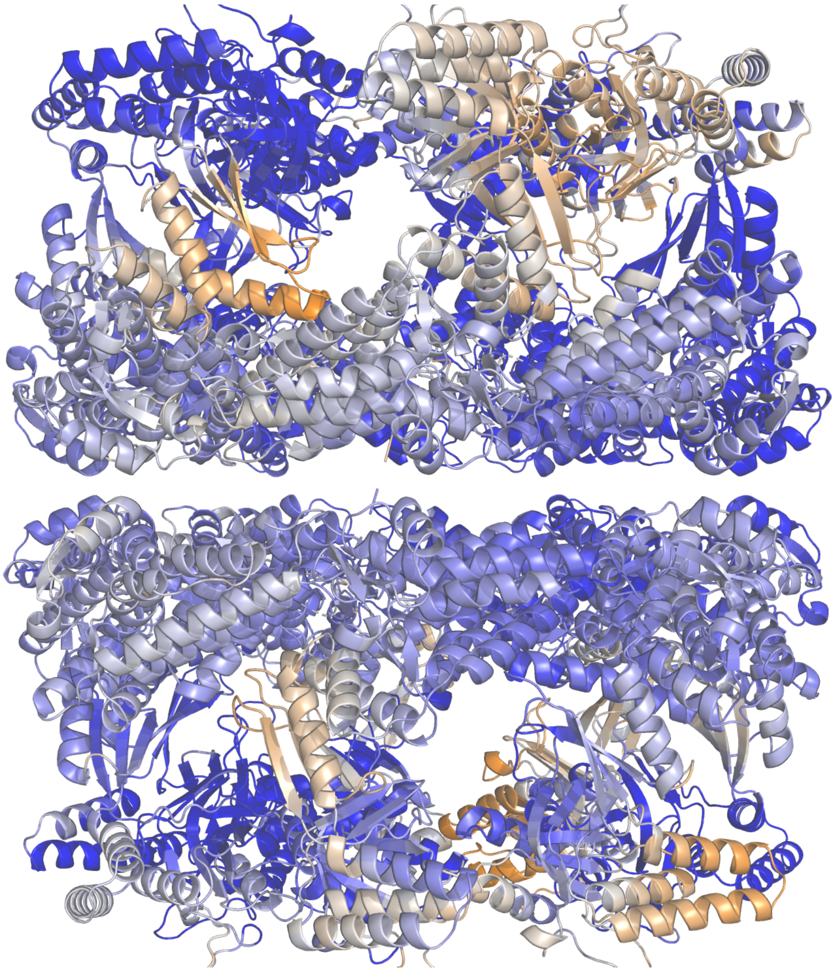

**Figure S6.** Average percentage change in DF score for each residue upon V72I mutation projected onto the starting structure of **D** ($P_{avg}$). The change is calculated from simulations of $\mathbf{D_{WT\text{-}AspAsp}}$ and $\mathbf{D_{V72I\text{-}AspAsp}}$. Colour intensities are codified in the colour bar shown in Figure 5 (main text).

#### RDF Plots from MD Simulations of $F_{WT}$ and $F_{V72I}$

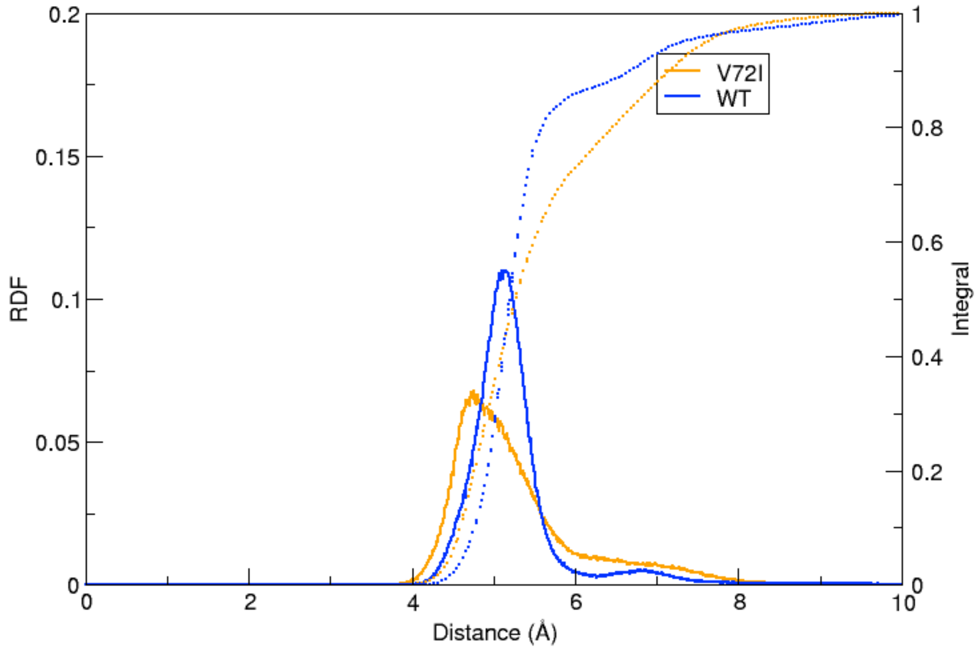

**Figure S7.** RDF plots of the Asp50:Cγ–Asp397:Cγ distance derived from MD metatrajectories of $\mathbf{F_{WT\text{-}AspAsp}}$ (blue) and $\mathbf{F_{V72I\text{-}AspAsp}}$ (orange). Dotted lines represent RDF integrals (right *y*-axis).

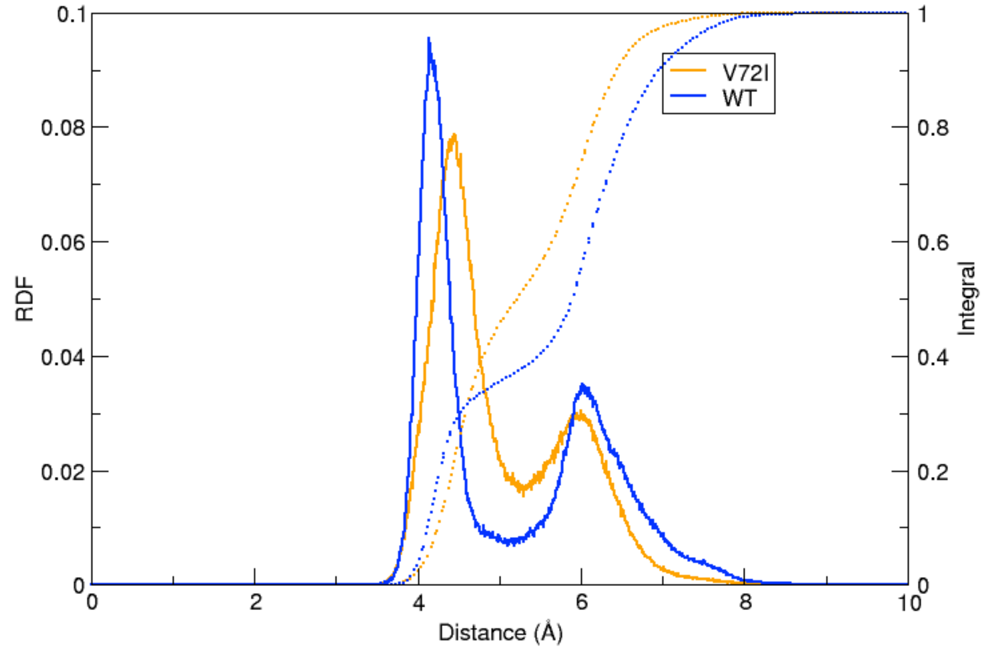

**Figure S8.** RDF plots of the Ash50:Cγ–Asp397:Cγ distance derived from MD metatrajectories of $\mathbf{F_{WT\text{-}AshAsp}}$ (blue) and $\mathbf{F_{V72I\text{-}AshAsp}}$ (orange). Dotted lines represent RDF integrals (right *y*-axis).

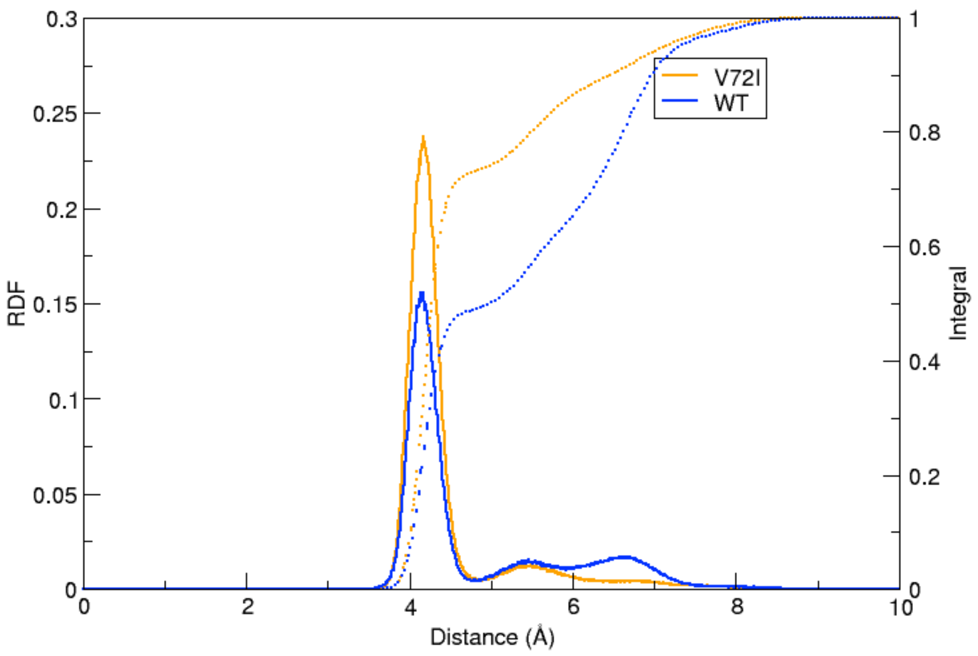

**Figure S9.** RDF plots of the Asp50:Cγ–Ash397:Cγ distance derived from MD metatrajectories of $\mathbf{F}_{\mathbf{WT\text{-}AspAsh}}$ (blue) and $\mathbf{F}_{\mathbf{V72I\text{-}AspAsh}}$ (orange). Dotted lines represent RDF integrals (right *y*-axis).

#### DF analysis on MD Simulations of $F_{WT}$ and $F_{V72I}$

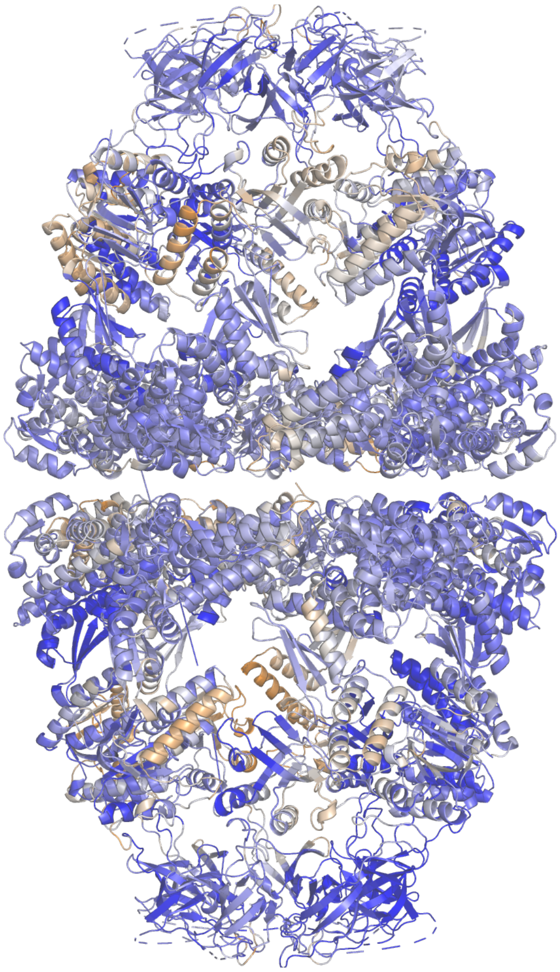

**Figure S10.** Average percentage change in DF score for each residue upon V72I mutation projected onto the starting structure of **F** ($P_{avg}$). The change is calculated from simulations of $\mathbf{F_{WT\text{-}AspAsp}}$ and $\mathbf{F_{V72I\text{-}AspAsp}}$. Colour intensities are codified in the colour bar shown in Figure 5 (main text).

#### Bibliography

1. D. A. Case, T. E. Cheatham, T. Darden, H. Gohlke, R. Luo, K. M. Merz, A. Onufriev, C. Simmerling, B. Wang and R. J. Woods, The Amber biomolecular simulation programs, *J. Comput. Chem.*, 2005, **26**, 1668-1688.
2. S. Miyamoto and P. A. Kollman, Settle: An analytical version of the SHAKE and RATTLE algorithm for rigid water models, *J. Comput. Chem.*, 1992, **13**, 952-962.
3. H. J. C. Berendsen, J. P. M. Postma, W. F. Van Gunsteren, A. DiNola and J. R. Haak, Molecular dynamics with coupling to an external bath, *J. Chem. Phys.*, 1984, **81**, 3684-3690.
4. R. Salomon-Ferrer, A. W. Götz, D. Poole, S. Le Grand and R. C. Walker, Routine Microsecond Molecular Dynamics Simulations with AMBER on GPUs. 2. Explicit Solvent Particle Mesh Ewald, *J. Chem. Theory Comput.*, 2013, **9**, 3878-3888.
5. R. J. Loncharich, B. R. Brooks and R. W. Pastor, Langevin dynamics of peptides: The frictional dependence of isomerization rates of N-acetylalanyl-N′-methylamide, *Biopolymers*, 1992, **32**, 523-535.
6. D. R. Roe and T. E. Cheatham, PTRAJ and CPPTRAJ: Software for Processing and Analysis of Molecular Dynamics Trajectory Data, *J. Chem. Theory Comput.*, 2013, **9**, 3084-3095.

7. S. Osuna, The challenge of predicting distal active site mutations in computational enzyme design, *Wiley Interdiscip. Rev.: Comput. Mol. Sci.*, 2021, **11**, e1502.
8. H. Shao, K. Oltion, T. Wu, J. E. Gestwicki, *Organic & Biomolecular Chemistry* 2020, **18**, 4157-4163
9. D. Saran, D. M. Held, D. H. Burke, *Nucleic Acids Research* 2006, **34**, 3201-3208
